# Antimicrobial resistance patterns reveal widespread multidrug-resistance among *Vibrio* species in shrimp hatchery environments of Bangladesh

**DOI:** 10.64898/2026.09.17.752394

**Authors:** Md Naimur Rahman, Shawon Ahmmed, Mohammad Shamsur Rahman

**Affiliations:** Aquatic Animal Health Group, Aquaculture Program, Department of Fisheries, University of Dhaka, Dhaka-1000, Bangladesh

**Keywords:** *Vibrio*, multidrug-resistant, shrimp hatchery, cross-resistance, co-selection, One Health

## Abstract

*Vibrio* species are ubiquitous in shrimp hatchery systems, where their opportunistic pathogenic characteristics threaten the vulnerable larval stage, and antibiotics are routinely applied for disease control and prevention. Therefore, studying *Vibrio* composition and its antibiotic resistance burden is critical for hatchery environments and public health. The present study characterized *Vibrio* composition and its antimicrobial resistance patterns in water-flow systems and postlarvae from shrimp hatcheries in Cox’s Bazar, Bangladesh. The findings revealed that *Vibrio* composition in these hatchery systems comprised nineteen different species. The Harveyi clade was the most dominant group, including *V. alginolyticus*, *V. rotiferianus*, and *V. harveyi*. The *Vibrio* isolates were highly resistant to erythromycin, ampicillin, and streptomycin, while remaining susceptible to chloramphenicol, nitrofurantoin, and gentamicin. Seven antibiotics, including trimethoprim-sulfamethoxazole, ampicillin, and erythromycin, were categorized as poorly active, suggesting their inefficacy against vibriosis outbreaks. More than 50% of the isolates had multiple antibiotic resistance index greater than 0.2 and 67.8% were categorized as multidrug-resistant. Both indices progressively increased along the water-flow systems and postlarvae had the highest values, pointing to antibiotic contamination within hatchery systems. Resistance gene screening identified *sul2* and *tetC* as most prevalent, while association analysis showed cross-resistance patterns among antibiotics, concordance between trimethoprim-sulfamethoxazole and streptomycin resistance with the presence of *sul2* and *strA*-*strB* genes, and co-selection of *sul2*, *ermB*, and *strA*-*strB* genes. The findings provide baseline data on *Vibrio* diversity and resistance burden, and inform antibiotic stewardship policies and monitoring strategies that support a One Health approach in shrimp hatchery systems.

**Importance:** Shrimp hatcheries supply the valuable seed that sustains shrimp aquaculture, providing jobs and income to many coastal communities. But hatchery environments involve highly intensified culture conditions and larvae are also susceptible to diseases like vibriosis. To manage diseases hatchery technicians heavily rely on antibiotics, used for both treatment and prevention. Many previous reports have mentioned the misuse and overuse of antibiotics in Bangladeshi shrimp hatcheries but data on antimicrobial resistance from this location remain limited. As aquatic environments are interconnected, there is a risk of antimicrobial resistance spreading from hatcheries to surrounding environments and beyond, posing a broader One Health concern that warrants immediate investigation. The significance of this study lies in characterizing *Vibrio* composition and phenotypic resistance in these hatcheries, providing diversity and resistance profiles that can help policymakers strengthen antibiotic regulations and guide hatchery technicians in developing practices that are aligned with the One Health framework.

## 1. Introduction

Shrimp culture is an important part of the advancing aquaculture industry, because of its fast growth, high market value, and strong global demand for seafood (Ayisi et al., 2017). Shrimp and prawns were the second most valuable traded species group (16%) among the aquatic animals in 2024 (Food and Agriculture Organization of the United Nations [FAO], 2026). Global shrimp exports reached 3.75 million tonnes in 2024, an increase of 2% over 2023, while imports were 3.71 million tonnes, valued at USD 25.4 billion (FAO, 2025). Among the shrimp exporting countries, Bangladesh produced 260,486 metric tons of shrimp and prawns and earned export revenue of USD 248.6 million in the 2023-2024 fiscal year (Department of Fisheries [DOF], 2024). This export value is largely driven by *Penaeus monodon*, the most cultured shrimp species in Bangladesh, accounting for 23.30% of total farm production (DOF, 2024), while it is the second most cultured shrimp species globally, accounting for approximately 12-13% of the global shrimp production in 2024 (FAO, 2026). This growing reliance on *Peanaeus monodon* culture requires a large amount of quality seed supply, which makes this industry highly hatchery-dependent. These highly demanded seeds are produced in the hatcheries of Cox’s Bazar, which contains around 47 *Penaeus monodon* hatcheries, making it the hub of shrimp seed supply in Bangladesh (DOF, 2024). The intensified culture conditions at the hatchery increase the risks of disease occurrence at this early life stage, resulting in high mortality and economic losses (Walker & Mohan, 2009). This vulnerability is due to the weak immunity system at the zoea to postlarvae stages, making shrimp highly susceptible to pathogenic infections (Vandenberghe et al., 1999). Susceptibility can also vary based on larval age and shrimp species. Zoea larvae are reported to be more susceptible than the later mysis stage and *Penaeus monodon* larvae may be more susceptible than *Macrobrachium rosenbergii* larvae (Prayitno & Latchford, 1995; Soto-Rodríguez et al., 2006).

Bacteria and viruses are considered the primary causative agents of shrimp larval diseases (Lightner & Redman, 1998). Among bacterial pathogens, opportunistic *Vibrio* species are a major cause of losses in penaeid hatcheries. The genus *Vibrio* is a group of Gram-negative rod shaped bacteria belonging to the Vibrionaceae family and Proteobacteria phylum, containing more than 100 known species (Onohuean et al., 2022; Thompson & Swings, 2006). They are also divided into several evolutionary clades including the Harveyi, Splendidus, Mediterranei, Orientalis, Nereis, Vulnificus, Diazotrophicus, Proteolyticus, and Fluvialis clades, among others (Jiang et al., 2021). Vibrios are naturally present in the water column, larval gut, body surface, and the live feed. They are also capable of causing disease under sub-optimal culture conditions (Decamp et al., 2008). Virulence in *Vibrio* species is strain-dependent, some strains can produce lethal exotoxins such as cysteine protease and haemolysins (Liu & Lee, 1999; Montero & Austin, 1999). These toxins are capable of damaging the mucosal barrier, allowing other bacteria to enter the internal organs of shrimp larvae (Soonthornchai et al., 2010). Among several diseases, a classical hatchery-related condition is luminous vibriosis, primarily caused by *V. harveyi* and *V. campbellii*, which are capable of causing up to 100% mortality in penaeid shrimp larvae and juveniles, with outbreaks reported across Asia, Australia, and Latin America (Kumar et al., 2021; Lio-Po, 2016). Another known disease associated with hatchery stage mass mortality is zoea II syndrome, caused by *V. alginolyticus*, *V. harveyi*, and *V. parahaemolyticus*, with incidence linked to prolonged stocking cycles and poor disinfection practices (Cuéllar-Anjel et al., 2010; Sathish Kumar et al., 2017). These bacteria are also reported to cause septic hepatopancreatic necrosis (SHPN), which affects postlarvae and juvenile stages of *Penaeus monodon* (Cuéllar-Anjel et al., 2010). *Vibrio* species are also known as zoonotic pathogens in humans, causing diarrheal illness and wound infections, and are transmitted through contaminated water and seafood (Actor, 2012). To control these emerging bacterial diseases, antibiotic usage has become an integrated part of shrimp aquaculture (Decamp et al., 2008; Kumar et al., 2021).

Shrimp aquaculture relies heavily on a broad spectrum of antibiotics and other antimicrobial agents such as parasiticides and heavy metals, with clinically relevant antibiotics reported to be used particularly in hatcheries (Thornber et al., 2020). The Food and Drug Administration (FDA) has authorized several antibiotics for use in aquaculture, including oxytetracycline, florfenicol, sarafloxacin, erythromycin, and sulphonamides combined with trimethoprim or ormethoprim (Serrano, 2005). Asian aquaculture has been reported to commonly use several antibiotics, including tetracycline, oxytetracycline, quinolones, trimethoprim, and sulphonamides, with three-quarter of major producing countries using oxytetracycline, sulphadiazine, and florfenicol (Lulijwa et al., 2020). In Bangladesh specifically, shrimp hatcheries of Cox’s Bazar have been reported to use up to 23 different antimicrobial products (Hinchliffe et al., 2018). Commonly used antibiotics included oxytetracycline, chlorotetracycline, amoxicillin, erythromycin, ciprofloxacin, co-trimoxazole, sulfadiazine, and sulfamethoxazole, with chloramphenicol used occasionally (Kawsar et al., 2026). These antibiotics were administered directly in the tank water containing postlarvae or brood shrimp (Aftabuddin et al., 2009). In these hatchery systems, antibiotics were applied both reactively, when disease signs were visible, and prophylactically for preventive purposes (Chowdhury et al., 2022). A significant proportion of antibiotics (70-90%) remains unmetabolized, which can accumulate in the aquaculture system or disseminate into the environment (Kawsar et al., 2026; Serrano, 2005). This constant selection pressure from antibiotics on the microbes has led to the emergence of antimicrobial-resistant (AMR) bacteria, specifically in *Vibrio* species as they are established pathogens in the hatchery systems (Bashar et al., 2026).

Antibiotics are among the most life-saving drugs in human and veterinary medicine, but the emergence of antimicrobial resistance has significantly reduced the efficacy of these treatments. This makes AMR a threat to global health in the current century (World Health Organization [WHO], 2017). Bacteria can develop antimicrobial resistance through several mechanisms, including enzymatic degradation or modification of antibiotics, extrusion by efflux pumps, target site modification, reduced permeability, bypass pathways, and biofilm formation (Afroze et al., 2025). Selection pressure from certain antibiotics can promote proliferation of bacteria that contains the selected resistant genes, favoring survival and spread within a population (Ferri et al., 2022). An alarming driver of the rapid dissemination of antimicrobial resistance among bacterial populations is horizontal gene transfer (HGT), which occurs via mobile genetic elements such as transposons and plasmids (Martínez et al., 2015). This process can be further escalated through co-selection, in which multiple resistance genes are carried on the same mobile genetic element. Separately, cross-resistance can increase the breadth of resistance, as a single mutation or acquired gene can result in resistance to multiple antibiotics of similar structure (Simjee et al., 2024). Antibiotic residues and resistant bacteria originating from shrimp hatchery systems can move easily between animals, the environment, and human populations, as aquatic environments are open and interconnected (Bashar et al., 2026). Furthermore, postlarvae produced in the hatchery systems are transferred to farms for the grow-out phase, which also poses a risk of disseminating AMR in farms and surrounding environments (Debnath et al., 2016). Ultimately, the AMR problem in a shrimp hatchery not only threatens the postlarvae themselves but also undermines the broader One Health goal of safeguarding interconnected human, animal, and environmental health (Kawsar, 2026). These problems underscore the need for regular monitoring of antimicrobial resistance in aquaculture systems, especially in the shrimp hatcheries, where antibiotics are often misused or overused.

There are several studies investigating antimicrobial resistance burden and resistance genes of *Vibrio* species in shrimp farms and market shrimp (Algammal et al., 2025; Bashar et al., 2026; Beshiru et al., 2020; Bourdonnais et al., 2024; Haque et al., 2023; Rahman et al., 2020; Sohidullah et al., 2025), but studies based on hatchery systems remain limited (Sotomayor et al., 2019; Yu et al., 2023). Additionally, several studies documented the overuse and misuse of antibiotics in Bangladeshi shrimp hatcheries (Bashar et al., 2026; Hinchliffe et al., 2018; Kawsar, 2026; Kawsar et al., 2026; Thornber et al., 2020), but investigation of resistance pattern and resistance genes is scarce. In order to address these gaps, we designed this study to systematically investigate *Vibrio* composition, resistance burden, and resistance genes. Samples were collected from the water-flow systems and postlarvae in shrimp hatcheries in Cox’s Bazar, Bangladesh. Results from antimicrobial susceptibility tests were used to classify the activity of antibiotics against *Vibrio* species. The multiple antibiotic resistance index (MAR) and prevalence of multidrug-resistant (MDR) isolates were also calculated to identify the high-risk source of contamination and evaluate the severity of resistance within shrimp hatchery systems. Furthermore, statistical analyses were performed to determine the association among phenotypic resistance patterns, between phenotypic resistance and resistance genes, as well as among resistance genes themselves in these hatchery systems. The findings will serve as baseline data for understanding broader marine *Vibrio* diversity and its current resistance burden in these hatchery systems. It will also provide essential guidelines on local antibiotic usage and implementation of antibiotic stewardship and monitoring strategies that support a One Health approach in shrimp hatchery systems.

## 2. Materials and methods

### 2.1. Sampling location

Cox’s Bazar district is the major *Penaeus monodon* postlarvae (PL) production area in Bangladesh. The PL are transported from here to the southern part of Bangladesh including Khulna, Bagerhat, and Satkhira districts for grow-out phase (Debnath et al., 2016). Cox’s Bazar district is considered a key area for establishing and operating hatcheries due to its favorable geography, which facilitates broodstock sourcing, high salinity and preferred water quality from the bay of Bengal. In this study, samples were collected from hatcheries in the two main upazilas for shrimp PL production, Cox’s Bazar Sadar (21.4367° N, 91.9933° E) and Ukhiya upazila (21.2450° N, 92.1375° E) (Figure 1), within Cox’s Bazar district.

**Figure 1.**
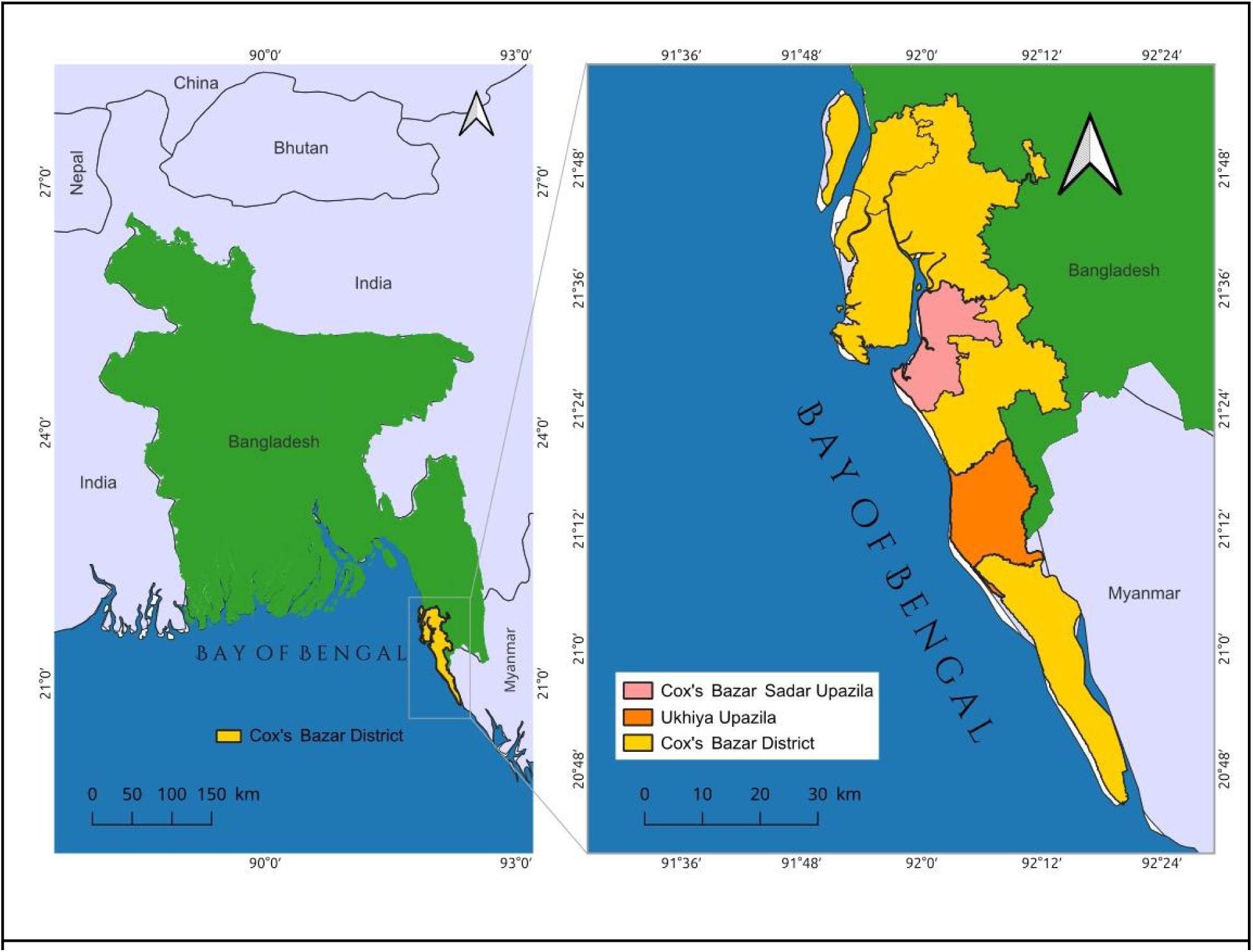
Map of the study area showing the sampling locations in Cox’s Bazar Sadar and Ukhiya upazila, Cox’s Bazar District, Bangladesh. The map was generated using QGIS v3.34.4.

### 2.2. Sample collection

Samples were collected during March to June of 2025. The collected samples included raw seawater (RSW), which was kept in the settlement tank for two days; treatment water (TW), which was collected after passing through the filtration steps (sand, coal, cartridge, and ultraviolet (UV)); rearing tank water (RTW), collected from the PL rearing tank; postlarvae (PL); and discarded water (DW) from the rearing tank outlet during water exchange or PL collection. The water samples were aseptically collected in a sterile bottle. Prior to sample collection, the bottle was washed three times with the source water. The PL samples were collected aseptically in a sterile zipper bag. Collected samples were packed with ice in an icebox and carried to the Microbiology Laboratory for further analysis. A total of 54 samples, including 39 water samples and 15 PL samples, were collected from seven hatcheries. Repeated samples were collected from several hatcheries when accessible during the sampling period, it was not on the purpose of fixed replicate or temporal schedule.

### 2.3. Sample preparation and bacterial isolation

Sample preparation was carried out following the protocol described in the Bacteriological Analytical Manual for *Vibrio* spp. isolation (Kaysner et al., 2004).The PL samples were washed three times with sterile saline water (0.85% NaCl) to get rid of the surface bacteria. Approximately 1g of PL sample was homogenized with 9 ml sterile saline water to prepare raw sample. The water and homogenized PL samples were serially diluted up to a 1000-fold dilution with sterile saline water. The samples were vortexed thoroughly before plating. A 100 µL aliquot of the undiluted and 100-fold diluted water samples, and the 10-fold and 1000 fold diluted PL samples, was spread onto Thiosulfate Citrate Bile Salts Sucrose (TCBS, Oxoid, UK) agar plates. In parallel, 1 ml of water samples or homogenized PL sample was added to separate tubes of 9 ml alkaline peptone water (APW, Oxoid, UK) and incubated for 6 hours at 37°C for *Vibrio* enrichment, which was streaked onto the TCBS agar plate. Both the spread and streaked TCBS plates were incubated at 37°C for 18 to 20 hours. Presumptive *Vibrio* colonies from TCBS spread plates of each sample were calculated as Colony Forming Units per mL (CFU/mL) using the following formula: CFU/mL = (Colony count) / (Dilution factor × Volume plated (mL)). Colony morphology characteristics on TCBS agar plates were recorded based on the protocol described by Breakwell et al. (2007). Colonies were selected based on sample source and morphological variation on TCBS agar, such as color, size, elevation, etc., and purified by streaking on TCBS and 2% NaCl added Tryptic Soy Agar (TSA, Oxoid, UK) plate. Selected colonies were stored in Luria-Bertani broth (LB, Oxoid, UK) with 30% glycerol supplementation at -80°C for further analysis. In this study, *Vibrio harveyi* JCM 33361^T^, *V. campbellii* JCM 1054^T^, *V. alginolyticus* HPV-03, *V. parahaemolyticus* BUVP-31 were used to study colony morphology and used as positive controls during PCR amplification. *Vibrio harveyi* JCM 33361^T^ and *V. campbellii* JCM 1054^T^ were procured from Japan Collection of Microorganisms (JCM), *V. alginolyticus* HPV-03 and *V. parahaemolyticus* BUVP-31 were previously identified through whole genome sequencing in our laboratory (data unpublished).

### 2.4. DNA extraction and molecular identification

Genomic DNA extraction was carried out by the methods described by Rahman et al. (2014). Freshly extracted DNA was quantified using NanoPhotometer® NP80 (Implen, Germany) and the high quality DNA (A260/280 ∼ 1.8) was stored at -20°C for further analysis. DNA from selected isolates (n = 59) was subjected to 16S rRNA gene amplification using the universal primers described by Weisburg et al. (1991), 27F (5′-AGAGTTTGATCCTGGCTCAG-3′) and 1492R (5′-CGGTTACCTTGTTACGACTT-3′) on a Biometra TAdvanced thermal cycler (Analytik Jena, Germany). The 25 µL reaction mixture comprised 12.5 µL GoTaq® G2 Green Master Mix (Promega, USA), 1 µL forward primer, 1 µL reverse primer, 9.5 µL nuclease-free water (Promega, USA) and 1 µL template DNA. Amplification conditions are given in Supplementary Table 1. Five microliters of PCR product was run on a runVIEW gel electrophoresis system (Cleaver Scientific, UK) using a 1% agarose gel containing 0.05 µL/mL ethidium bromide (EtBr, SRL, India) in 1X Tris-acetate-EDTA buffer (TAE, Thermo Scientific, USA) at 100 V for 40 minutes, alongside both 100 bp and 1 kb ladders (Promega, USA). The gel was visualized on a UVP PhotoDoc-It imaging system (Analytik Jena, Germany) to confirm the presence of ∼1500 bp bands. The PCR products were sent to BDGenomes (Bangladesh) for Sanger sequencing (Sanger et al., 1977), which was performed by Genecreate Biotech (China). Both the forward and reverse reads were visualized, manually trimmed, and aligned using the Unipro UGENE v53.1 (Okonechnikov et al., 2012), with ambiguous base calls resolved by comparison against BLAST search results. The resulting consensus sequences were subjected to homology search using BLASTn (http://blast.ncbi.nlm.nih.gov) against the GenBank database (Altschul et al., 1990; Altschul et al., 1997).

### 2.5. Phylogenetic tree construction

Consensus sequences of the 16S rRNA gene were aligned along with the type strain sequences of each identified species, obtained from the NCBI nucleotide database, using MAFFT v7.526 in auto mode (Katoh & Standley, 2013). The multiple sequence alignment was visualized through AliView v1.3 (Larsson, 2014) and trimmed using ClipKIT v2.12 with the smart-gap option (Steenwyk et al., 2020). Two sequences, *Vibrio coralliilyticus* msr39 and *Vibrio xuii* msr46 were excluded from tree construction due to their insufficient sequence length. A maximum likelihood phylogenetic tree was constructed using IQ-TREE v3.1.2 (Wong et al., 2026), with model selection performed using ModelFinder (Kalyaanamoorthy et al., 2017) and branch support assessed via 1000 ultrafast bootstrap replicates (Hoang et al., 2018) and 1000 SH-aLRT replicates. The sequence of *Photobacterium leiognathi* ATCC 25521T (NR_115541) was used to root the tree and as an outgroup. The resulting tree was visualized using ggtree v4.3.0 (Yu et al., 2018) in R v4.6.1 (R Core Team, 2026).

### 2.6. Antimicrobial susceptibility testing

Antimicrobial susceptibility of the selected *Vibrio* isolates (n = 59) was tested by Kirby-Bauer disc diffusion method as described in CLSI guideline M02-A11 (Performance Standards for Antimicrobial Disk Susceptibility Tests) and M45 (Methods for Antimicrobial Dilution and Disk Susceptibility Testing of Infrequently Isolated or Fastidious Bacteria) (Clinical & Laboratory Standards Institute [CLSI], 2012, 2015). Briefly, freshly grown colonies were used to prepare inoculum suspension in sterile saline water (0.85% NaCl), and turbidity was adjusted to match the 0.5 McFarland standard (HiMedia, India). A sterile wooden-stick cotton swab was used to spread inoculum suspension on the Mueller-Hinton agar (MHA, Oxoid, UK), by rotating the plate three times at a 60° angle. The plate was left to dry for 5-10 minutes, after which the antibiotic discs were placed on the MHA. A total of 24 antibiotics (Oxoid, UK) from 11 antimicrobial classes were used in this study (Table 1). The agar plates were incubated at 37°C for 18 to 20 hours in an inverted position. The diameter of the antibiotic inhibition zone was measured to the nearest whole millimeter. The measured diameter was used to interpret the antibiotic susceptibility profile as susceptible, intermediate, and resistant according to the criteria of CLSI guideline M45 for *Vibrio* spp. (CLSI, 2015). The zone interpretation criteria of azithromycin, ceftriaxone, erythromycin, nitrofurantoin, nalidixic acid, and streptomycin were not available in the M45 guideline, so they were interpreted using CLSI guideline M100 (CLSI, 2023).

**Table 1.**
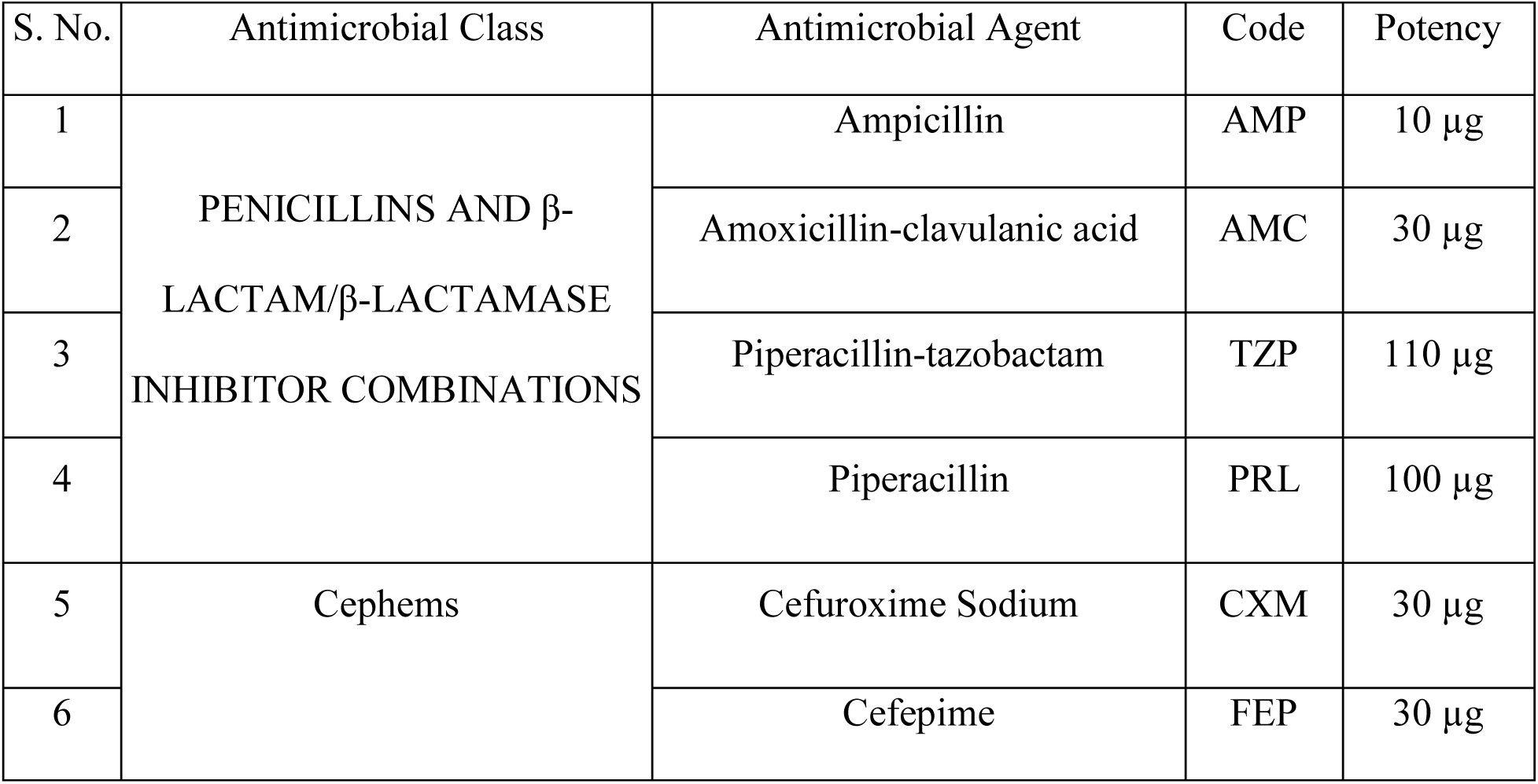
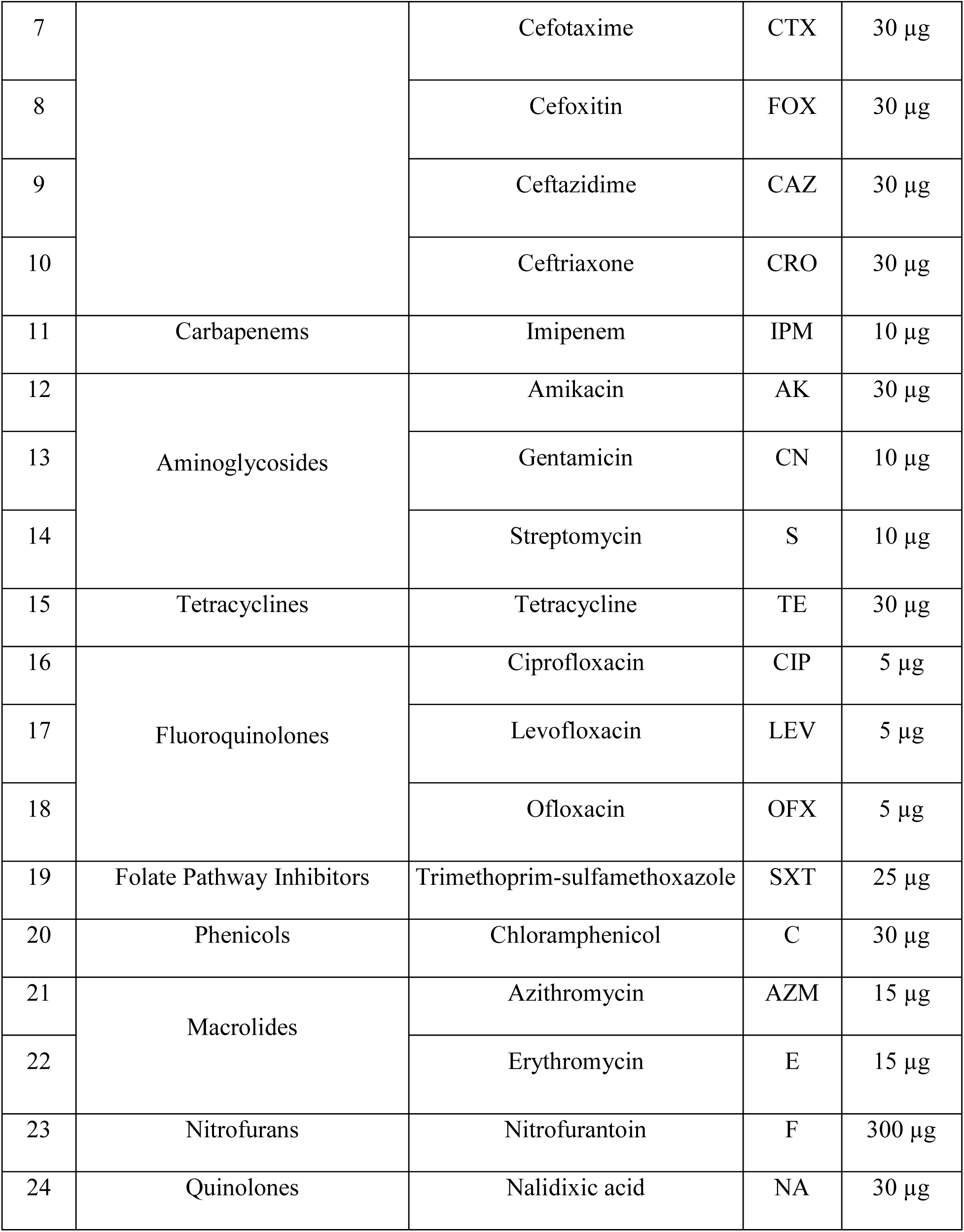
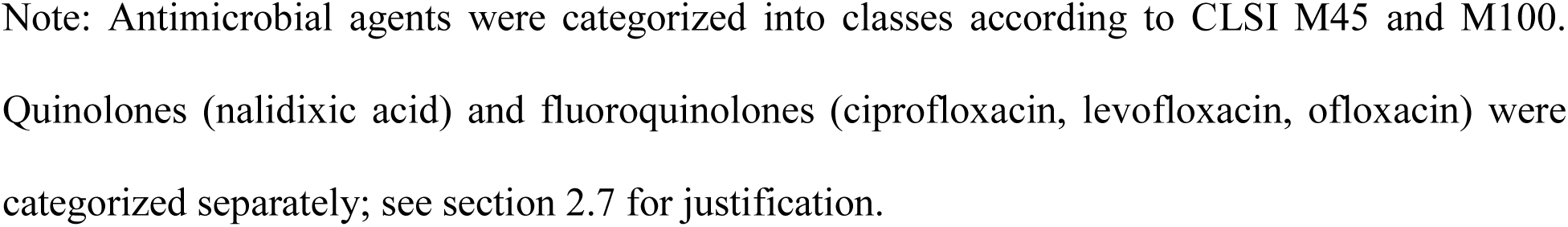
List of antimicrobial agents and their class, code, and potency used in this study.

| S. No. | Antimicrobial Class | Antimicrobial Agent | Code | Potency |
| --- | --- | --- | --- | --- |
| 1 | PENICILLINS AND β-LACTAM/β-LACTAMASE INHIBITOR COMBINATIONS | Ampicillin | AMP | 10 μg |
| 2 | | Amoxicillin-clavulanic acid | AMC | 30 μg |
| 3 | | Piperacillin-tazobactam | TZP | 110 μg |
| 4 | | Piperacillin | PRL | 100 μg |
| 5 | Cephems | Cefuroxime Sodium | CXM | 30 μg |
| 6 | | Cefepime | FEP | 30 μg |
| 7 |  | Cefotaxime | CTX | 30 µg |
| 8 |  | Cefoxitin | FOX | 30 µg |
| 9 |  | Ceftazidime | CAZ | 30 µg |
| 10 |  | Ceftriaxone | CRO | 30 µg |
| 11 | Carbapenems | Imipenem | IPM | 10 µg |
| 12 | Aminoglycosides | Amikacin | AK | 30 µg |
| 13 |  | Gentamicin | CN | 10 µg |
| 14 |  | Streptomycin | S | 10 µg |
| 15 | Tetracyclines | Tetracycline | TE | 30 µg |
| 16 | Fluoroquinolones | Ciprofloxacin | CIP | 5 µg |
| 17 |  | Levofloxacin | LEV | 5 µg |
| 18 |  | Ofloxacin | OFX | 5 µg |
| 19 | Folate Pathway Inhibitors | Trimethoprim-sulfamethoxazole | SXT | 25 µg |
| 20 | Phenicol | Chloramphenicol | C | 30 µg |
| 21 | Macrolides | Azithromycin | AZM | 15 µg |
| 22 |  | Erythromycin | E | 15 µg |
| 23 | Nitrofurans | Nitrofurantoin | F | 300 µg |
| 24 | Quinolones | Nalidixic acid | NA | 30 µg |

Note: Antimicrobial agents were categorized into classes according to CLSI M45 and M100. Quinolones (nalidixic acid) and fluoroquinolones (ciprofloxacin, levofloxacin, ofloxacin) were categorized separately; see section 2.7 for justification.

### 2.7. Antibiotic resistance characterization indexing analysis

Multiple antibiotic resistance (MAR) index was calculated by dividing the total number of antibiotics an isolate is resistant to by the total number of antibiotics tested per isolate (Krumperman, 1983). A MAR index value greater than 0.2 indicates that the organism is from an environment with high antibiotic usage, which may facilitate development of antibiotic resistance (Titilawo et al., 2015). Additionally, antibiotic resistance pattern abundance (ARPA) indicates the variability of antibiotic resistance patterns in an environment; higher values imply higher variability. ARPA was calculated by the formula described by Deng et al. (2020), as ARPA = (number of resistance types / total number of isolates).

Variation coefficient percentage (VC%) was used to determine the variability of antibiotic activity against the *Vibrio* isolates and it was calculated by the formula, VC% = (100 × SD/Mean of the zone of inhibition diameters) (Lakhssassi et al., 2005). For each antibiotic, activity score was independently assigned based on susceptibility percentage (S%) and based on variation coefficient percentage (VC%). Activity score 0 was assigned to S% < 25 or VC% ≥ 100, 1 to S% 25–50 or VC% 75–100, 2 to S% and VC% both 50–75, 3 to S% 75–100 or VC% 25–50, and 4 to S% = 100 or VC% < 25. The two scores were then summed to obtain a final activity score (0–8) for each antibiotic, which was classified as inactive (0), poorly active (≥1 to <5), fairly active (≥5 to <8), or very active (8) (Sony et al., 2021). An isolate was defined as multi-drug resistant (MDR) when it was found resistant to at least one antibiotic from three or more antimicrobial classes (Magiorakos et al., 2012). For MDR assignment, the antibiotics were categorized into different classes based on CLSI M45, supplemented with CLSI M100. Nalidixic acid and fluoroquinolones were treated as separate classes, due to their distinct chemical structures and antimicrobial spectra (Fàbrega et al., 2008).

### 2.8. Molecular characterization of antimicrobial resistance genes

All the selected isolates (n = 59) were screened for five antimicrobial resistance genes through polymerase chain reaction (PCR). The target genes included *bla*_TEM_, *ermB*, *strA-strB*, *sul2*, and *tetC*, which confer resistance to beta-lactams, erythromycin, streptomycin, sulphonamide, and tetracycline, respectively (Table 2). The PCR reaction mixture and gel electrophoresis procedures were as previously described in the “DNA extraction and molecular identification” section. The PCR amplification conditions are listed in the Supplementary Table 1.

**Table 2.**
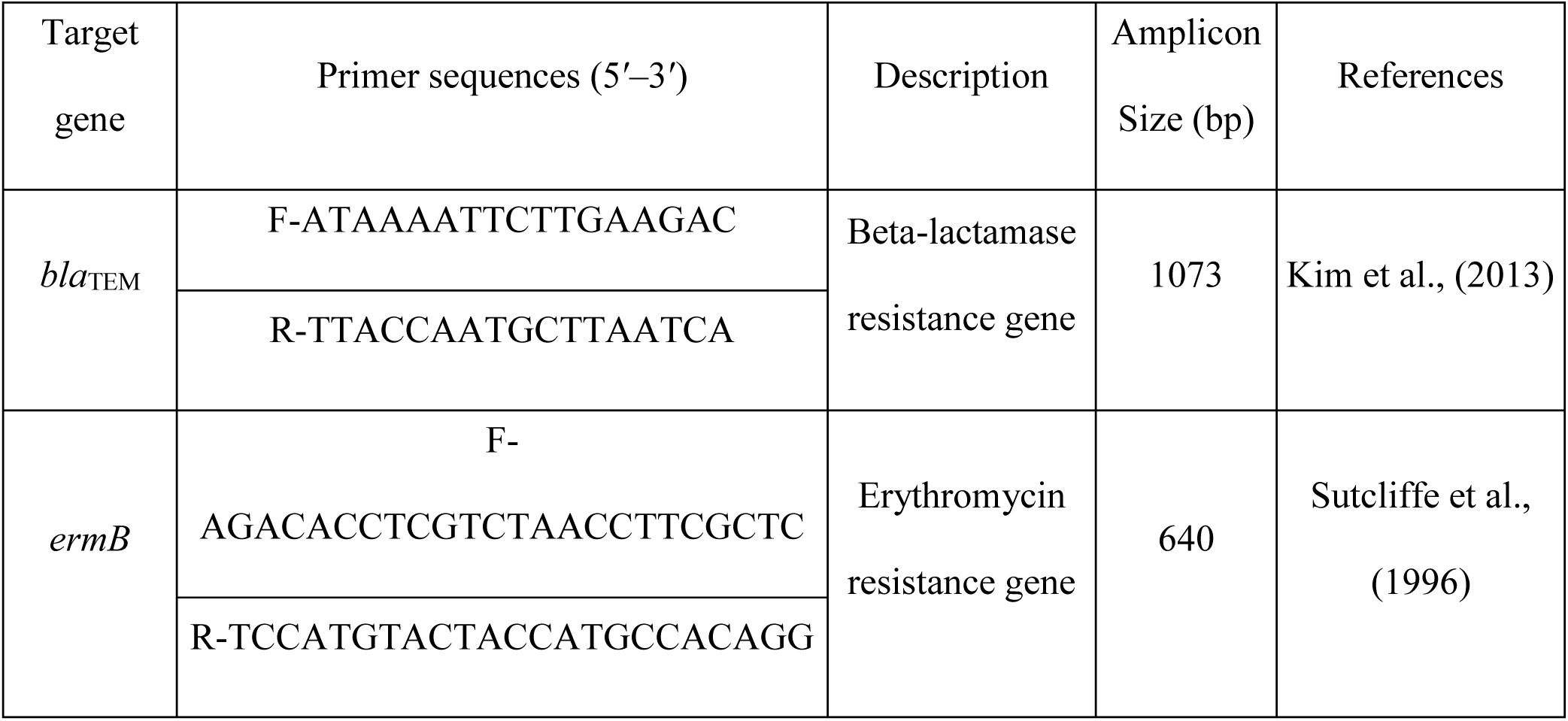
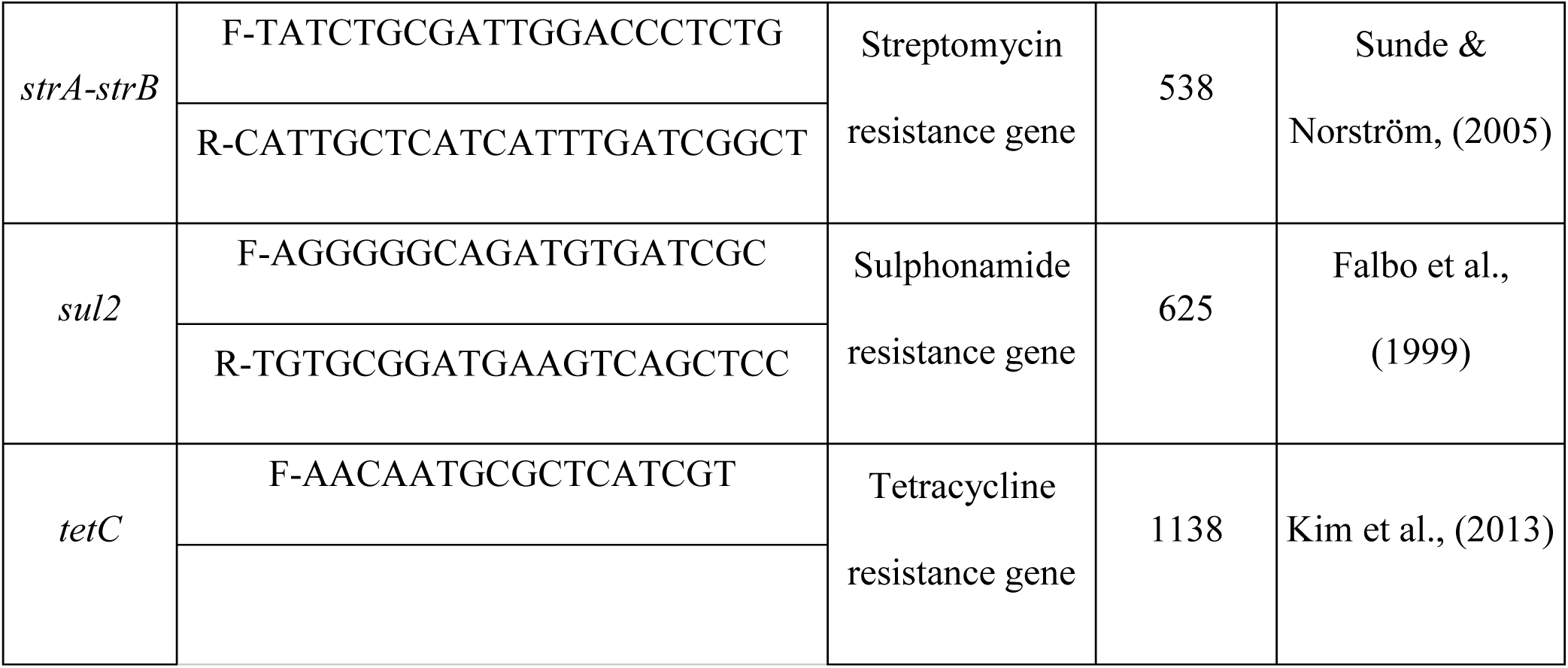
Lists of antimicrobial resistance genes screened in this study.

### 2.9. Statistical analysis

Statistical analyses were performed following the recommendations provided by Olsen (2014). Presumptive *Vibrio* count (CFU/mL) was log_10_-transformed for statistical analysis and visualization. Normality of raw data and model residuals were assessed using the Shapiro-Wilk test and Q-Q plots, homogeneity of variance was assessed using Levene’s test. For multiple group comparisons of presumptive *Vibrio* count (log_10_CFU/mL) across sources, Welch’s ANOVA followed by Games-Howell post hoc test was used as the model residual distribution was normal but variance homogeneity was violated. For multiple group comparisons of MAR indexes across sources, Kruskal-Wallis test followed by Dunn’s post-hoc pairwise comparisons with Holm-Šídák correction was used as both model residuals and variance of homogeneity were borderline and sample sizes per sources were small and unequal (n = 8-19). For comparison of MAR index between two locations, Wilcoxon rank sum test was used as data were not normally distributed. Correlations between presumptive *Vibrio* count (log_10_CFU/mL) and MAR index were assessed using Spearman’s rank correlation analysis due to data normality violation and non-linear relationship. The joint relationship between presumptive *Vibrio* count, MAR index, and MDR status was assessed using multiple logistic regression with Firth’s penalized likelihood, applied due to near-separation observed in the standard maximum-likelihood model (Firth, 1993). Associations among antimicrobial phenotypic resistance, between phenotypic resistance and isolation source, between antimicrobial resistance genes and antimicrobial phenotypic resistance, and among antimicrobial resistance genes themselves were assessed using Fisher’s exact test. Fisher’s exact test was chosen over Chi-square tests due to small expected cell counts in contingency tables. Resulting *P* values from Fisher’s exact test for multiple comparisons were adjusted using Benjamini-Hochberg false discovery rate (FDR) method. A *P* value less than 0.05 was considered statistically significant; where correction for multiple comparison was applied, an adjusted *P* value (*P*_adj_) less than 0.05 was considered statistically significant. All statistical analyses were performed with car v3.1.5 (Fox & Weisberg, 2019), logistf v1.26.1 (Heinze et al., 2025), and rstatix v1.1.0 (Kassambara, 2026) packages, and data visualization was performed using ggplot2 v4.0.3 (Wickham, 2016) and ComplexHeatmap v2.28.0 (Gu et al., 2016) in R v4.6.1 (R Core Team, 2026).

## 3. Results

### 3.1 Presumptive *Vibrio* spp. count and morphological variations

Presumptive *Vibrio* spp. colonies from the TCBS spread plates of each sample were recorded and presented as log_10_-transformed values of CFU/mL (Figure 2). The colony count did not differ significantly among the water sources (RSW, TW, RTW, DW), notably, even the treated water (TW) (Games-Howell test following Welch’s ANOVA, *P*_adj_ > 0.05). The PL samples had significantly higher colony count than the water sources (*P*_adj_ < 0.001). A total of 158 colonies were preliminarily selected based on their morphological characteristics on TCBS agar plates. After excluding duplicate or repetitive colonies from the same sample, a total of 59 colonies were finally selected for further study (Figure 3). Morphological variations were recorded based on seven traits, viz., color, shape, size, margin, elevation, texture, and opacity. Observed color variations were green, yellow, and cream. Most of the isolates were circular shaped with an entire margin. Green and yellow isolates had mostly moderate size and smooth or smooth and glossy texture, whereas cream color isolates had large size and mucoid texture. Green and cream color isolates had convex elevation, whereas yellow colonies had umbonate, flat, and convex elevation. Almost all of them were opaque.

**Figure 2.**
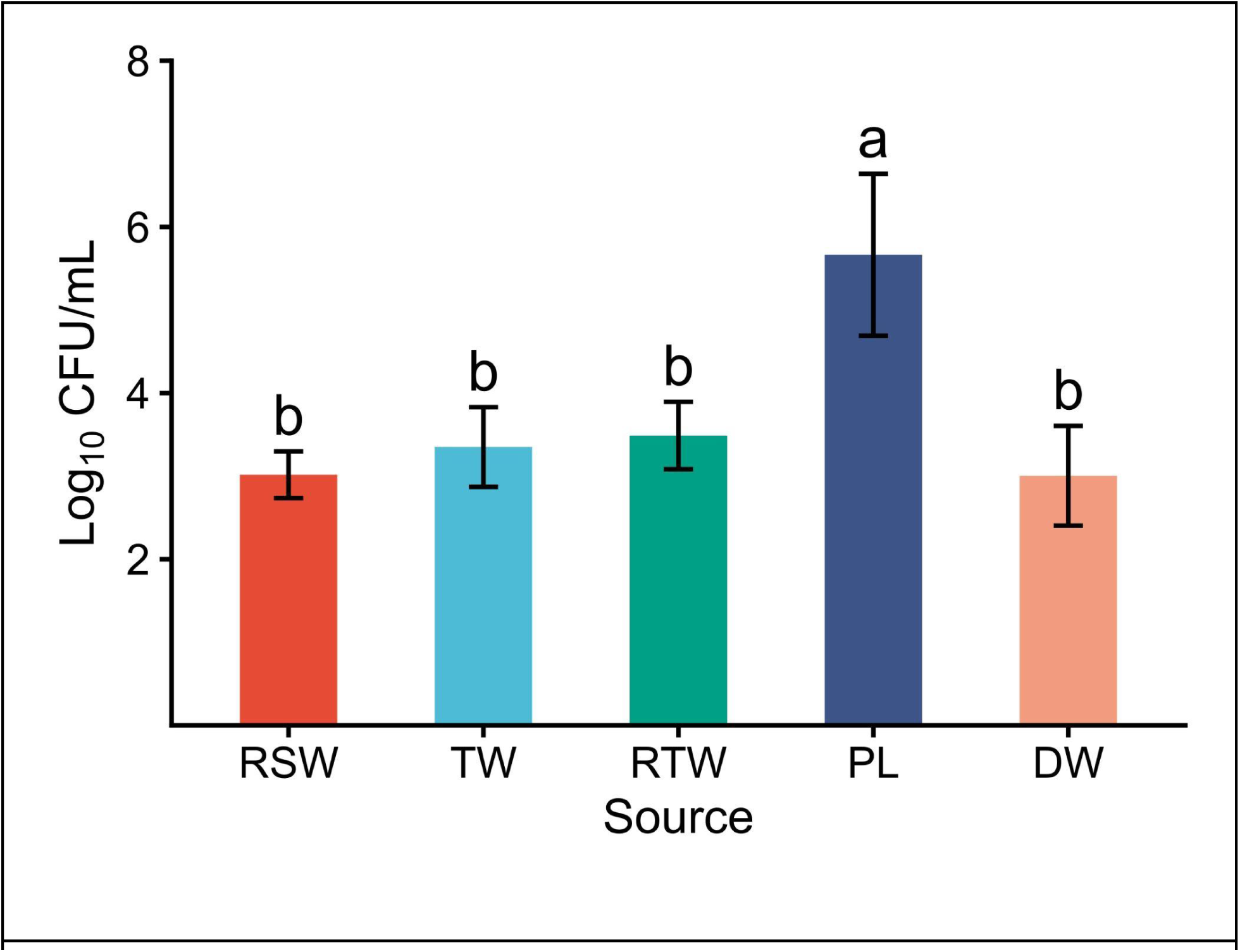
Presumptive *Vibrio* spp. count (Log_10_CFU/mL) is presented as a bar chart (mean ± sd) across sources. Mean values not sharing letters are significantly different (Games-Howell post hoc, *P* < 0.05). Overall, differences between sources were assessed by Welch’s ANOVA test (F(4, 15.1) = 22.2, *P* < 0.001), followed by Games-Howell post hoc comparisons (all significant pairs, *P* < 0.001). Abbreviations: RSW, raw seawater; TW, treated water; RTW, rearing tank water; PL, postlarvae; DW, discarded water.

**Figure 3.**
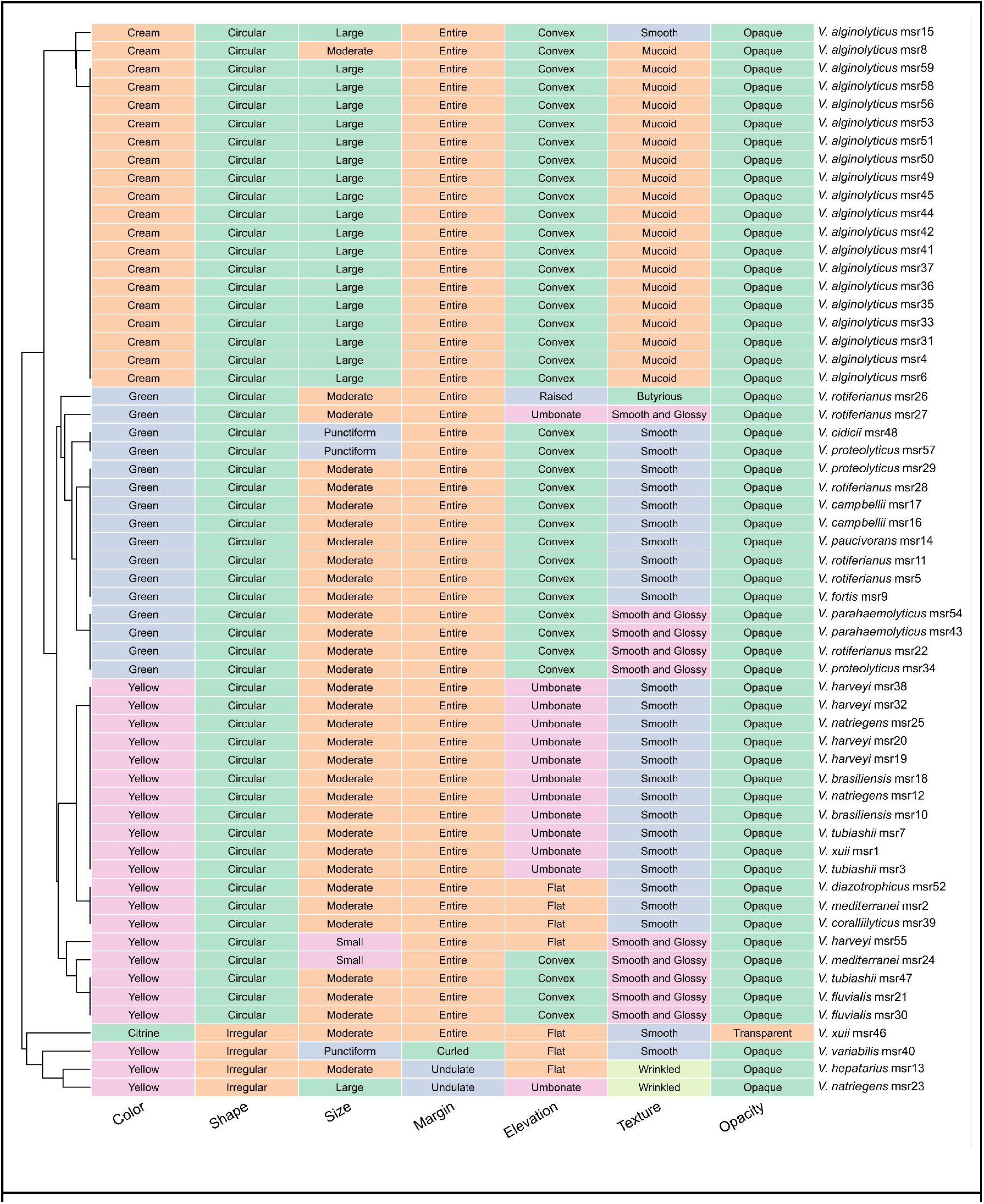
Morphological variations of the selected isolates (n = 59) on the TCBS plates are presented as a heatmap. Size variations were categorized based on colony diameters as punctiform (<0.5 mm), small (0.5 to 1 mm), moderate (1 to 3 mm), and large (>3 mm). Morphology similarity between isolates was assessed using Hamming distance (proportion of discordant traits across 7 morphological characteristics) and hierarchically clustered using average linkage (UPGMA).

### 3.2. Molecular identification by 16S rRNA gene sequencing

The selected isolates (n = 59) were identified by 16S rRNA gene sequencing and homology searching on the BLASTn program of the NCBI. The species identity was given based on the top BLAST hit, ranked by maximum bit score and lowest E-value (Supplementary Table 2). A maximum likelihood (ML) phylogenetic tree was created using IQ-TREE to determine the taxonomic placement and evolutionary relatedness of the isolates along with their corresponding type strain sequences retrieved from the NCBI nucleotide database (Figure 4). The isolates of the same species and their type strains were closely clustered together in the ML tree, confirming their species-level identification. The tree captured 19 diversified *Vibrio* species and they were divided into nine clades based on existing literature: the Harveyi clade (*V. natriegens*, *V. alginolyticus*, *V. parahaemolyticus*, *V. harveyi*, *V. rotiferianus*, and *V. campbellii*), Proteolyticus clade (*V. proteolyticus*), Splendidus clade (*V. fortis*), Mediterranei clade (*V. mediterranei* and *V. variabilis*), Nereis clade (*V. brasiliensis* and *V. xuii*), Orientalis clade (*V. tubiashii* and *V. hepatarius*), Diazotrophicus clade (*V. diazotrophicus*), Vulnificus clade (*V. cidicii*), and Fluvialis clade (*V. fluvialis*). *V. paucivorans* clustered separately from these established clades. The members of each clade clustered close to each other. The clade isolates had nearly zero branch length within their own species cluster, showing low intraspecific genetic diversity at the 16S rRNA locus among the hatchery isolates.

**Figure 4.**
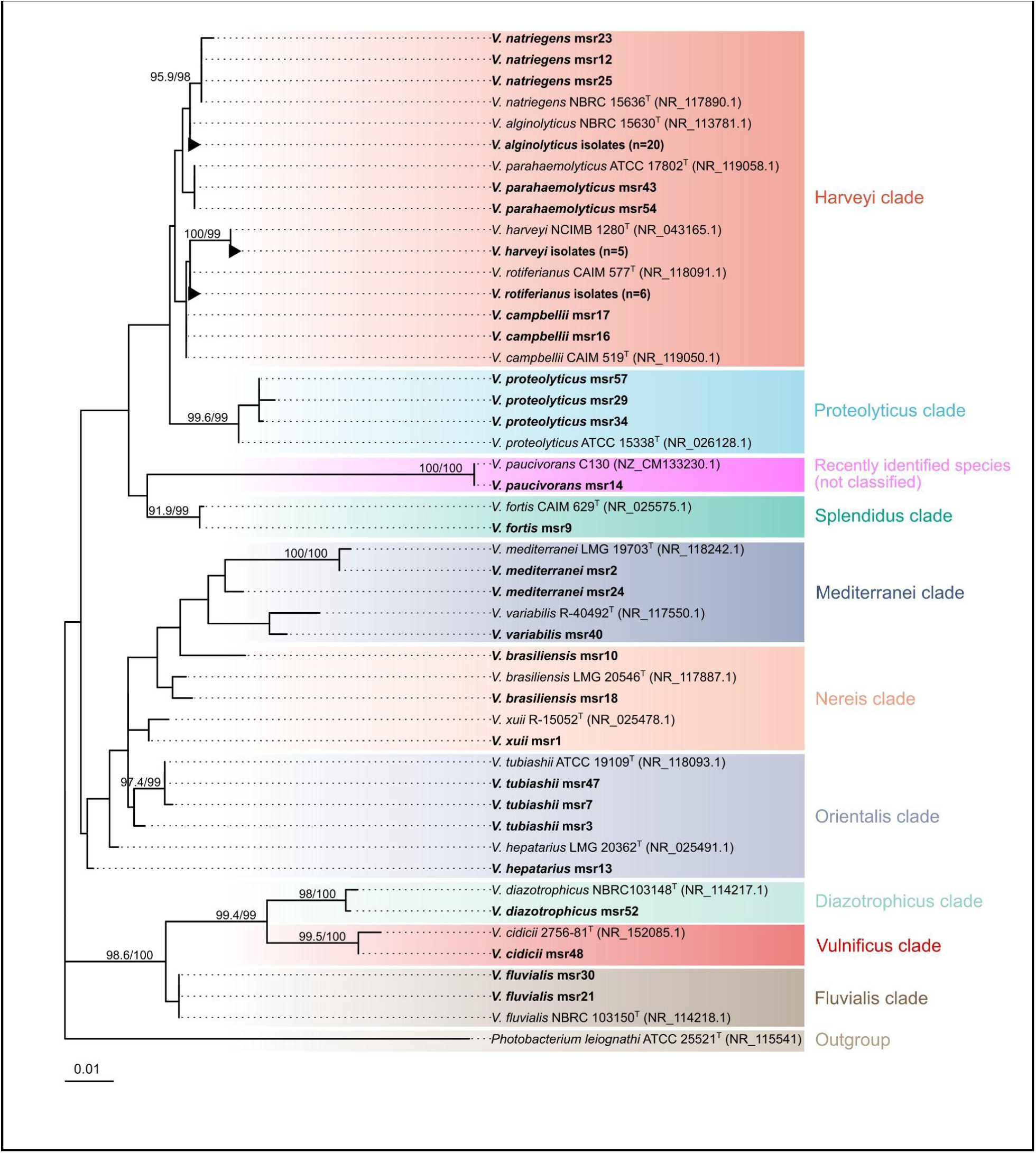
Phylogenetic tree of *Vibrio* isolates and type strain sequences of each identified species, based on 16S rRNA gene sequences, constructed using Maximum Likelihood analysis. *Photobacterium leiognathi* ATCC 25521^T^ (NR_115541) was used as an outgroup. Node values represent SH-aLRT/ultrafast bootstrap support (only values ≥80/95 are shown). The scale bar represents 0.01 substitutions per nucleotide site. The clade marking was done based on existing literature. The clade colors are based on species color gradients. *Vibrio coralliilyticus* LV_118 and *Vibrio xuii* LV_134 were excluded from tree construction due to their short sequence length.

### 3.3. *Vibrio* species diversity within hatchery system

A total of 59 isolates were selected from two locations (Cox’s Bazar Sadar and Ukhiya Upazila) and identified to the species level using 16S rRNA gene sequencing. The isolates were obtained from five sources, viz., raw seawater (RSW), treated water (TW), rearing tank water (RTW), postlarvae (PL), and discarded water (DW). All identified isolates belonged to the genus *Vibrio*, comprising 19 different species (Figure 5). *V. alginolyticus* was the most dominant species (n = 20), and was recovered from all sources. The next most frequently isolated species were *V. rotiferianus* (n=6), *V. harveyi* (n=5), *V. tubiashii* (n=3), *V. proteolyticus* (n=3), and *V. natriegens* (n=3). The highest species diversity was found in RSW and PL samples, each comprising 8 different species. Additionally, *V. alginolyticus*, *V. rotiferianus*, *V. natriegens*, and *V. fluvialis*were shared between RTW and PL samples. Overall, *V. alginolyticus* was the only species recovered from all sources, suggesting widespread distribution in the hatchery system, from raw seawater through the rearing process to postlarvae and final discharge.

**Figure 5.**
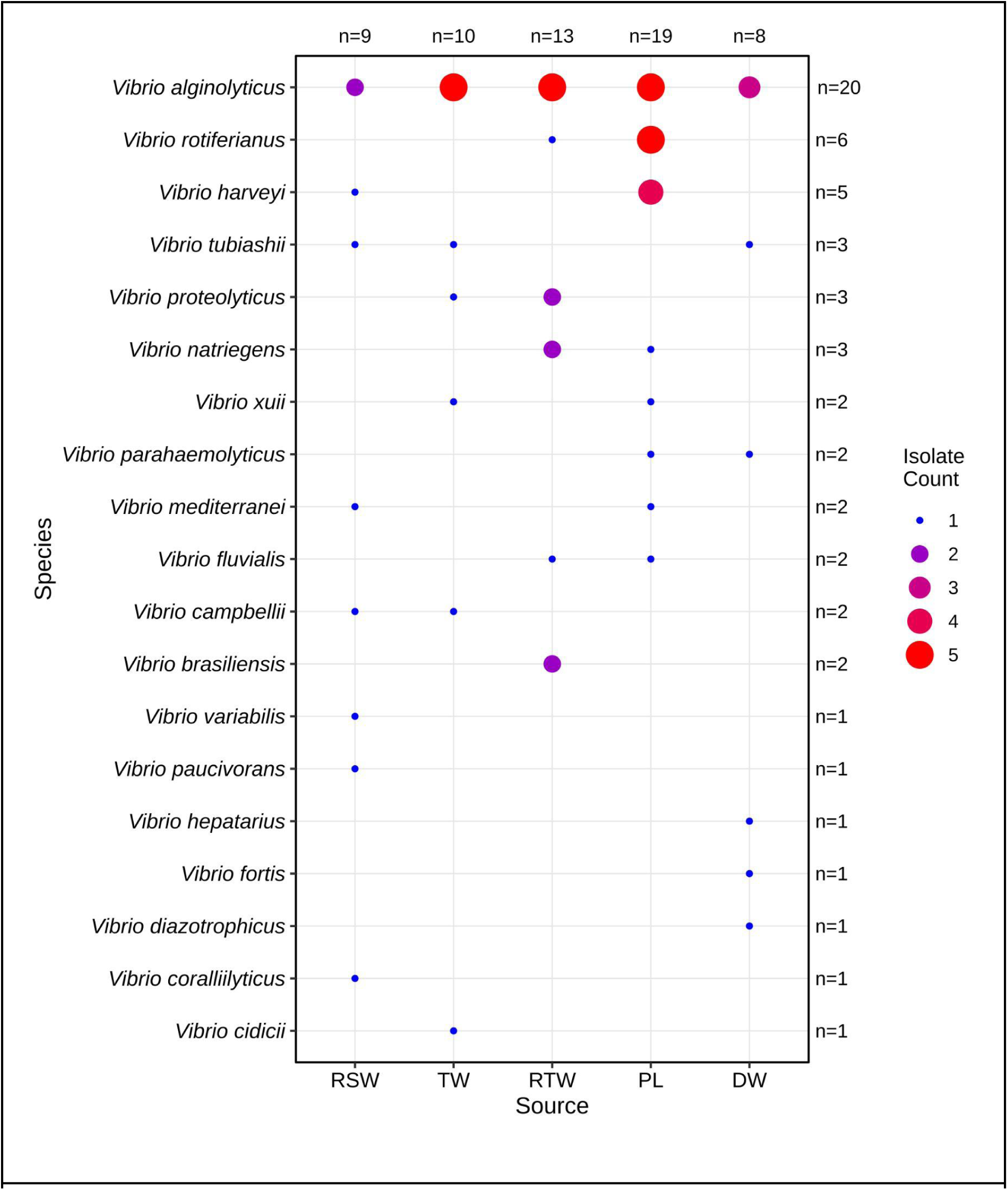
*Vibrio* species (n = 59) distribution and prevalences across sampling sources are presented as a dot plot. The dot size and color intensity presents the isolate number per species-source combination. The numbers at the top present the isolate number per source, and numbers at the right present total isolates per species. Abbreviations: RSW: raw seawater; TW: treated water; RTW: rearing tank water; PL: postlarvae; DW: discarded water.

The correspondence between morphological traits of the isolates and their molecular identification was inconsistent (Figure 3). The morphological traits of the identified *V. alginolyticus* isolates were similar and predictive of species identity as they clustered together in the heatmap. Whereas, green colonies had very similar traits and clustered together, yet they were identified as different species, including *V. rotiferianus*, *V. campbellii*, *V. proteolyticus*, and *V. parahaemolyticus*. The yellow colored colonies also had limited variations, yet identified as several species, including *V. harveyi*, *V. natriegens*, *V. tubiashii*, and *V. brasiliensis*.

### 3.4. Antimicrobial resistance phenotypes

In total, 59 *Vibrio* isolates were subjected to antimicrobial susceptibility testing against 24 antibiotics from 11 antimicrobial classes using the Kirby-Bauer disc diffusion method (Table 1). The zone of inhibition diameters were used to calculate the variation coefficient percentage (VC%), which revealed that ampicillin (AMP) exhibited the highest variable activity (VC% = 125.8%) against the *Vibrio* isolates, indicating highly unpredictable in vivo antibacterial action. Higher variability (VC% > 25%) was also observed for E (70%), SXT (66.2%), S (63%), NA (61%), AZM (57.7%), OFX (46.2%), CIP (44.6%), LEV (43.6%), CXM (39.4%), PRL (37.1%), and TE (34.6%). Low variability (VC% ≤ 25%) was observed for C (20.8%), AMC (18.6%), CAZ (17.8%), FOX (16.6%), F (14.6%), FEP (14%), TZP (13.6%), CTX (13.2%), CRO (12%), IPM (11.6%), AK (10.6%), and CN (7.2%) (Figure 6A).

**Figure 6.**
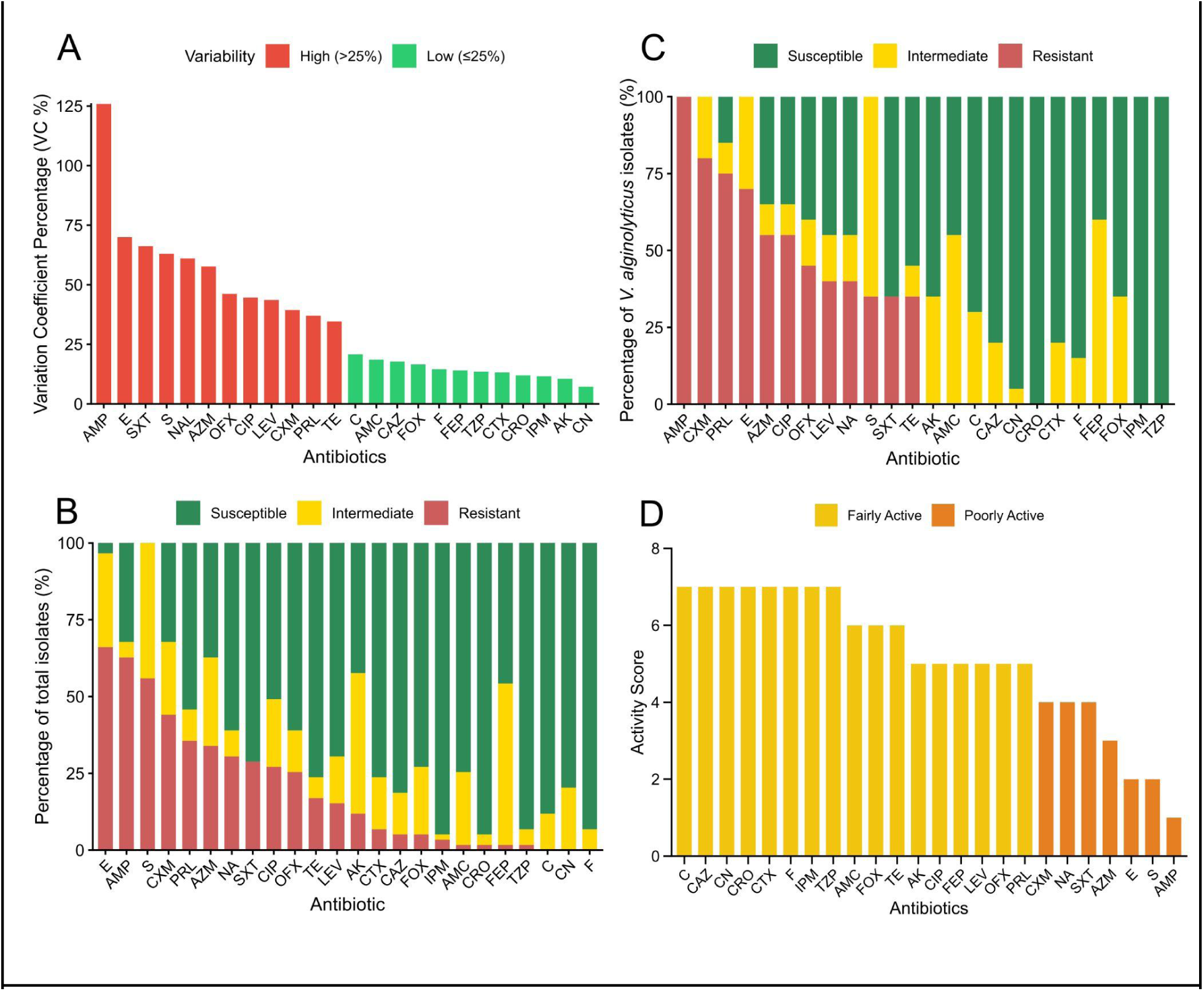
Antibiotics resistance patterns, variation coefficient percentage (VC%), and activity scores of the antibiotics. (A) Variation coefficient percentage (VC%) of the antibiotics are presented as bar plots.(B) Percentages of all isolates (n = 59) that are susceptible, intermediate, or resistant to each antibiotic presented as bar plots. A VC% greater than 25% was considered high variation (red color) and VC% lower than 25% was considered as low variation (green color). (C) Percentages of *V. alginolyticus* isolates (n = 20) that are susceptible, intermediate, or resistant to each antibiotic presented as bar plots. (D) Activity scores of the antibiotics are presented as bar plots. They were categorized as fairly active with score (≥5 to <8), and poorly active with score (≥1 to <5). Abbreviations: AMP: ampicillin, AMC: amoxicillin-clavulanic acid, TZP: piperacillin-tazobactam, PRL: piperacillin, CXM: cefuroxime sodium, FEP: cefepime, CTX: cefotaxime, FOX: cefoxitin, CAZ: ceftazidime, CRO: ceftriaxone, IPM: imipenem, AK: amikacin, CN: gentamicin, S: streptomycin, TE: tetracycline, CIP: ciprofloxacin, LEV: levofloxacin, OFX: ofloxacin, SXT: trimethoprim-sulfamethoxazole, C: chloramphenicol, AZM: azithromycin, E: erythromycin, F: nitrofurantoin, NA: nalidixic acid

The isolates showed varied resistance patterns to the antibiotics tested. Two isolates (3.4%) were not resistant to any antibiotics, 20 isolates (33.9%) showed low resistance (1–3 antibiotics), 23 isolates (39%) showed moderate resistance (4–6 antibiotics), and 14 isolates (23.7%) showed high resistance (7–12 antibiotics) (Table 3). The *Vibrio* isolates showed high resistance (>50%) to E (66.1%), AMP (62.7%), and S (55.9%). Moderate resistance (10%–50%) was seen in CXM (44.1%), PRL (35.6%), AZM (33.9%), NA (30.5%), SXT (28.8%), CIP (27.1%), OFX (25.4%), TE (16.9%), LEV (15.3%), and AK (11.9%). Low resistance (<10%) was seen in CTX (6.8%), CAZ (5.1%), FOX (5.1%), IPM (3.4%), AMC (1.7%), CRO (1.7%), FEP (1.7%), and TZP (1.7%). No isolates were found resistant to C, CN, and F (Figure 6B). At species level, *V. alginolyticus* (n = 20) showed high resistance to AMP (100%), CXM (80%), PRL (75%), E (70%), AZM (55%), and CIP (55%), whereas remained fully susceptible (100%) to CRO, IPM, TZP (Figure 6C).

**Table 3.**
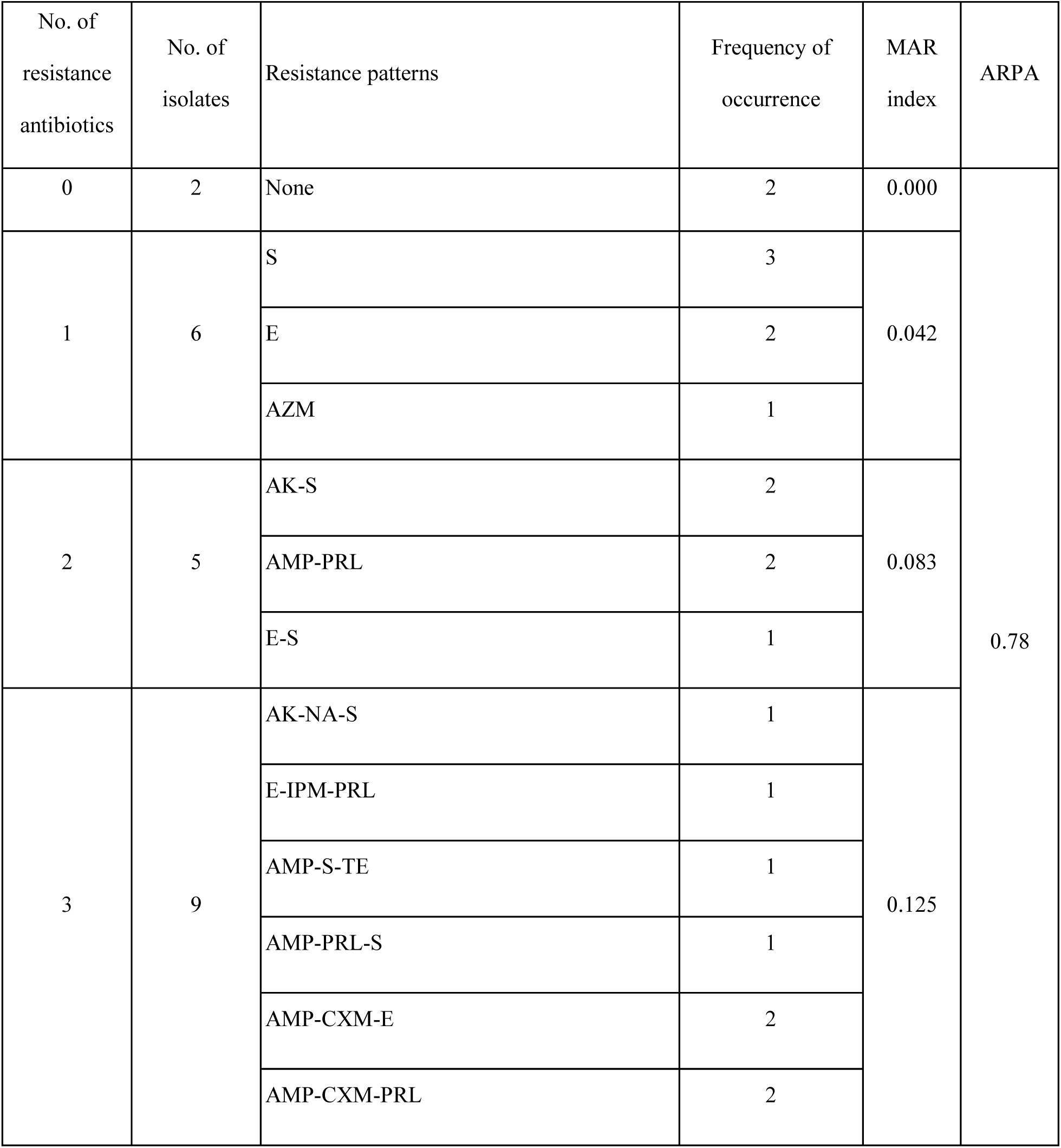
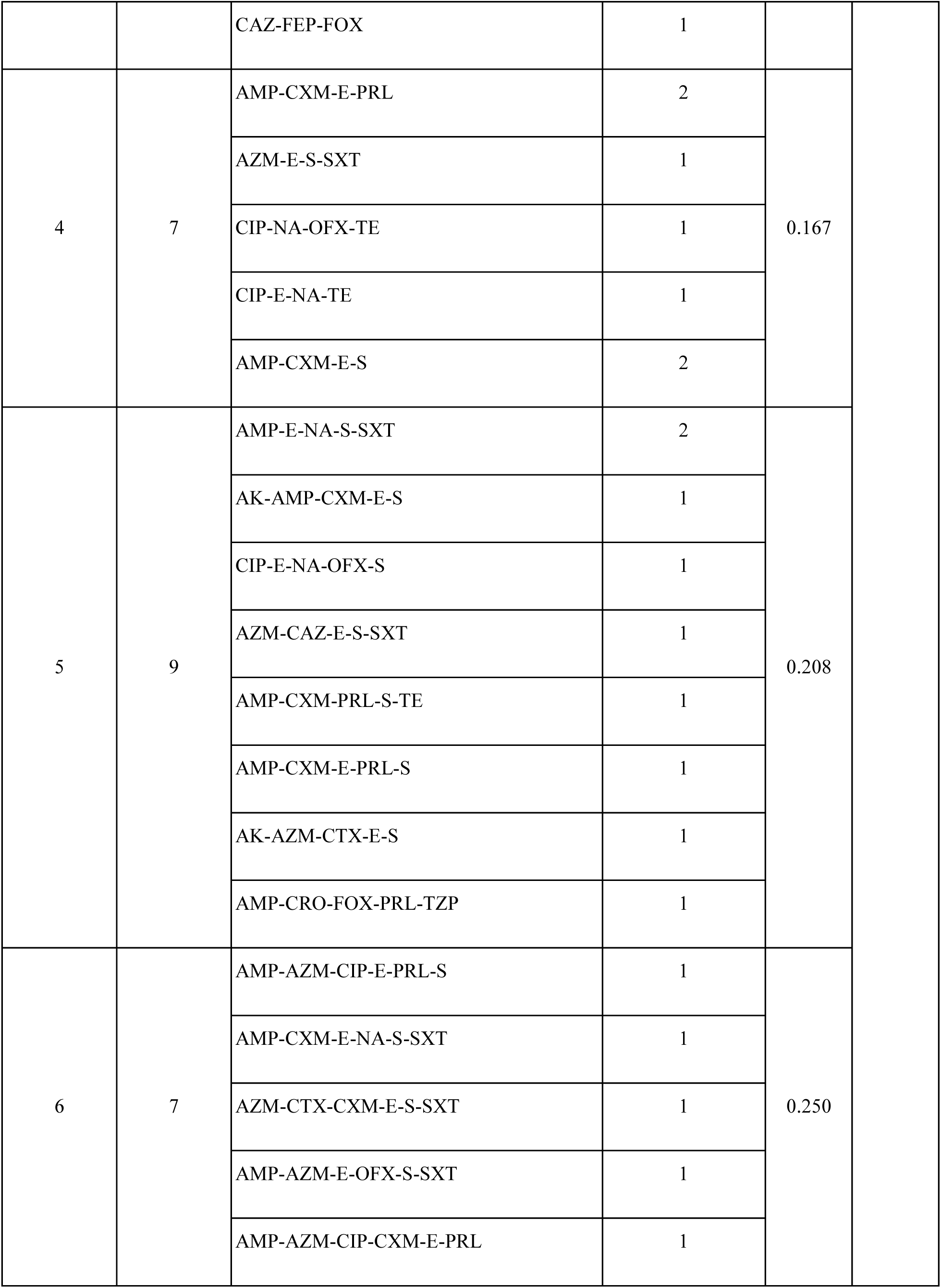
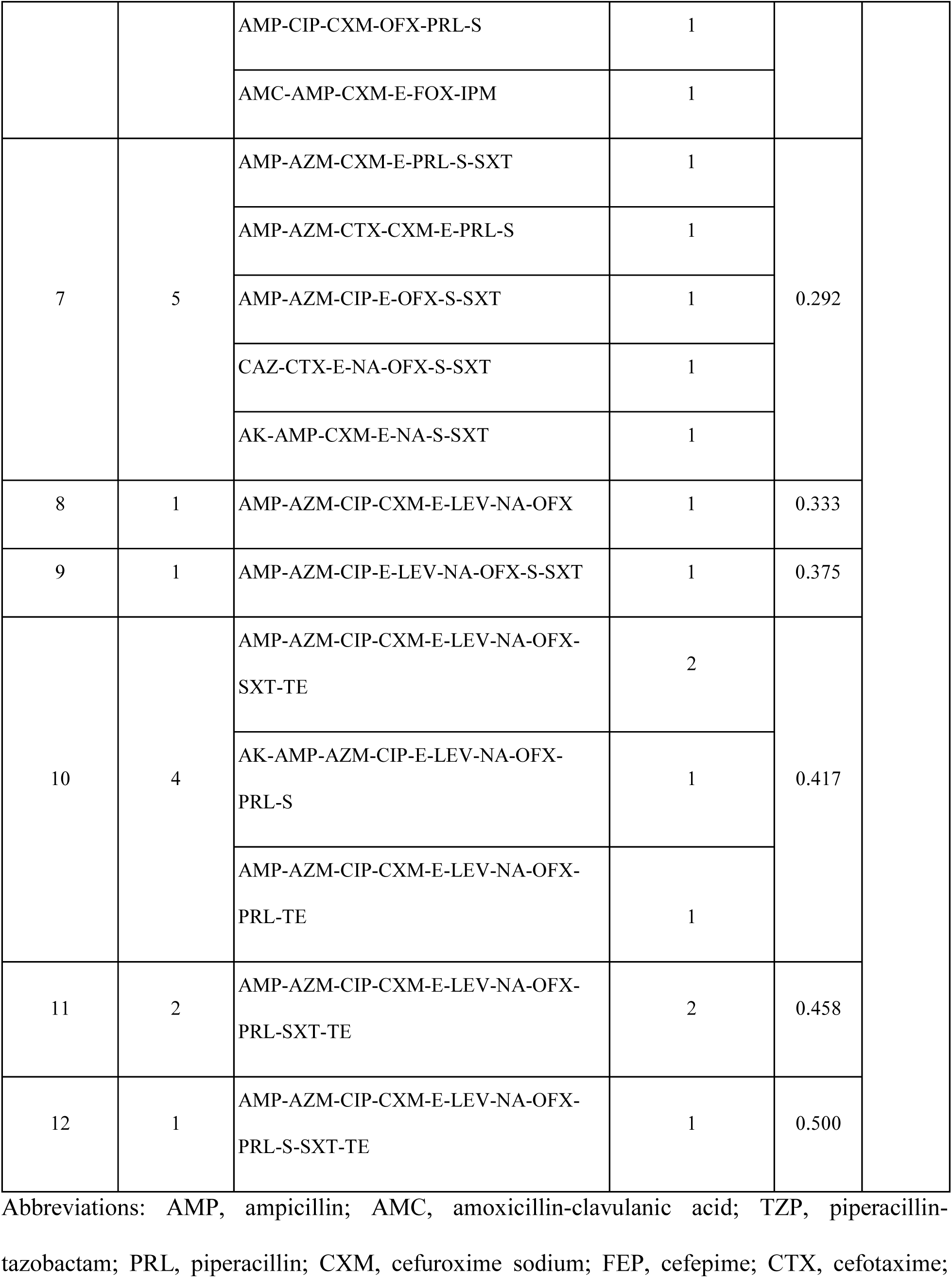
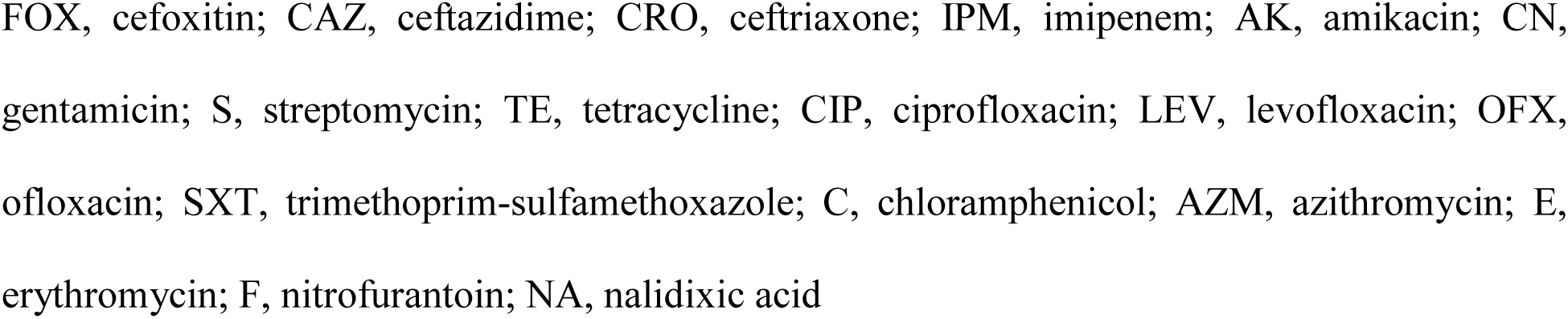
Pattern of multiple antibiotic resistance phenotypes and index of the *Vibrio* species.

| No. of resistance antibiotics | No. of isolates | Resistance patterns | Frequency of occurrence | MAR index | ARPA |
| --- | --- | --- | --- | --- | --- |
| 0 | 2 | None | 2 | 0.000 | 0.78 |
| 1 | 6 | S | 3 | 0.042 |  |
|  |  | E | 2 |  |  |
|  |  | AZM | 1 |  |  |
| 2 | 5 | AK-S | 2 | 0.083 |  |
|  |  | AMP-PRL | 2 |  |  |
|  |  | E-S | 1 |  |  |
| 3 | 9 | AK-NA-S | 1 | 0.125 |  |
|  |  | E-IPM-PRL | 1 |  |  |
|  |  | AMP-S-TE | 1 |  |  |
|  |  | AMP-PRL-S | 1 |  |  |
|  |  | AMP-CXM-E | 2 |  |  |
|  |  | AMP-CXM-PRL | 2 |  |  |
|  |  | CAZ-FEP-FOX | 1 |  |  |
| 4 | 7 | AMP-CXM-E-PRL | 2 | 0.167 |  |
|  |  | AZM-E-S-SXT | 1 |  |  |
|  |  | CIP-NA-OFX-TE | 1 |  |  |
|  |  | CIP-E-NA-TE | 1 |  |  |
|  |  | AMP-CXM-E-S | 2 |  |  |
| 5 | 9 | AMP-E-NA-S-SXT | 2 | 0.208 |  |
|  |  | AK-AMP-CXM-E-S | 1 |  |  |
|  |  | CIP-E-NA-OFX-S | 1 |  |  |
|  |  | AZM-CAZ-E-S-SXT | 1 |  |  |
|  |  | AMP-CXM-PRL-S-TE | 1 |  |  |
|  |  | AMP-CXM-E-PRL-S | 1 |  |  |
|  |  | AK-AZM-CTX-E-S | 1 |  |  |
|  |  | AMP-CRO-FOX-PRL-TZP | 1 |  |  |
| 6 | 7 | AMP-AZM-CIP-E-PRL-S | 1 | 0.250 |  |
|  |  | AMP-CXM-E-NA-S-SXT | 1 |  |  |
|  |  | AZM-CTX-CXM-E-S-SXT | 1 |  |  |
|  |  | AMP-AZM-E-OFX-S-SXT | 1 |  |  |
|  |  | AMP-AZM-CIP-CXM-E-PRL | 1 |  |  |
|  |  | AMP-CIP-CXM-OFX-PRL-S | 1 |  |  |
|  |  | AMC-AMP-CXM-E-FOX-IPM | 1 |  |  |
| 7 | 5 | AMP-AZM-CXM-E-PRL-S-SXT | 1 | 0.292 |  |
|  |  | AMP-AZM-CTX-CXM-E-PRL-S | 1 |  |  |
|  |  | AMP-AZM-CIP-E-OFX-S-SXT | 1 |  |  |
|  |  | CAZ-CTX-E-NA-OFX-S-SXT | 1 |  |  |
|  |  | AK-AMP-CXM-E-NA-S-SXT | 1 |  |  |
| 8 | 1 | AMP-AZM-CIP-CXM-E-LEV-NA-OFX | 1 | 0.333 |  |
| 9 | 1 | AMP-AZM-CIP-E-LEV-NA-OFX-S-SXT | 1 | 0.375 |  |
| 10 | 4 | AMP-AZM-CIP-CXM-E-LEV-NA-OFX-SXT-TE | 2 | 0.417 |  |
|  |  | AK-AMP-AZM-CIP-E-LEV-NA-OFX-PRL-S | 1 |  |  |
|  |  | AMP-AZM-CIP-CXM-E-LEV-NA-OFX-PRL-TE | 1 |  |  |
| 11 | 2 | AMP-AZM-CIP-CXM-E-LEV-NA-OFX-PRL-SXT-TE | 2 | 0.458 |  |
| 12 | 1 | AMP-AZM-CIP-CXM-E-LEV-NA-OFX-PRL-S-SXT-TE | 1 | 0.500 |  |

Abbreviations: AMP, ampicillin; AMC, amoxicillin-clavulanic acid; TZP, piperacillin- tazobactam; PRL, piperacillin; CXM, cefuroxime sodium; FEP, cefepime; CTX, cefotaxime;
FOX, cefoxitin; CAZ, ceftazidime; CRO, ceftriaxone; IPM, imipenem; AK, amikacin; CN, gentamicin; S, streptomycin; TE, tetracycline; CIP, ciprofloxacin; LEV, levofloxacin; OFX, ofloxacin; SXT, trimethoprim-sulfamethoxazole; C, chloramphenicol; AZM, azithromycin; E, erythromycin; F, nitrofurantoin; NA, nalidixic acid

### 3.5. Activity scoring of antibiotics

Final activity scores were assigned based on the percentage of susceptible isolates and the variation coefficient (VC%) (Figure 6D). Based on these scores, 17 antibiotics, viz., C, CAZ, CN, CRO, CTX, F, IPM, TZP, AMC, FOX, TE, AK, CIP, FEP, LEV, OFX, and PRL were categorized as fairly active (activity score 5–7). Seven antibiotics, viz., CXM, NA, SXT, AZM, E, S, and AMP, were categorized as poorly active (activity score 1–4). AMP had the lowest activity score of 1. None of the antibiotics were classified as very active or inactive against the *Vibrio* isolates.

### 3.6. Indexes and pattern of multiple antibiotic resistance phenotypes

MAR index was calculated for the 59 *Vibrio* isolates across all sources. MAR index ranged from 0 to 0.5, with 30 (50.8%) isolates having greater than 0.2 values with a median of 0.21 (Figure 7A). At species level, *V. alginolyticus*, *V. rotiferianus*, and *V. harveyi* accounted for 76.7% (n = 23) of the 30 isolates exceeding MAR threshold of greater than 0.2. The MAR index of most highly resistant isolates are listed in Table 4. Among them, five isolates were *V. alginolyticus* and one *V. rotiferianus*. The highest MAR index (0.5) isolate, *V. alginolyticus* msr8 showed resistance to 12 antibiotics, viz., AMP, PRL, CXM, S, TE, CIP, LEV, OFX, SXT, AZM, E, and NA. The top most resistant isolates showed a common resistance pattern to seven antibiotics including AMP, CIP, LEV, OFX, AZM, E, NA. Although, *V. alginolyticus* msr36 and *V. alginolyticus* msr56 were isolated from different sources, and locations they showed the same resistant pattern. The MAR index of *Vibrio* isolates differed significantly across sources (Kruskal-Wallis test, χ² = 9.64, df = 4, *P* = 0.047), and increased progressively among water sources from RSW to DW (median = 0.13 to 0.21), and PL isolates had a higher MAR index (median = 0.25) than the water isolates (Figure 7B). Following Dunn’s post-hoc analysis with Holm correction, there were only significant differences between RSW and PL samples (*P*_adj_ = 0.03), other pairs were statistically non-significant (*P*_adj_ > 0.05). MAR indexes of the isolates from Ukhiya and Cox’s Bazar Sadar did not show significant differences (Wilcoxon rank sum test, W = 322.5, *P* = 0.72) (Figure 7B). Antibiotics resistance pattern abundance (ARPA) was also calculated for all isolates, revealing 46 resistance patterns and ARPA value of 0.78 (Table 3). The highest seen resistant pattern (3x) was resistant to streptomycin (S), the rest combinations were unique, eleven patterns were seen two times each, and the rest were seen one time each. At species level, *V. alginolyticus* isolates (n = 20) had an ARPA of 0.8.

**Figure 7.**
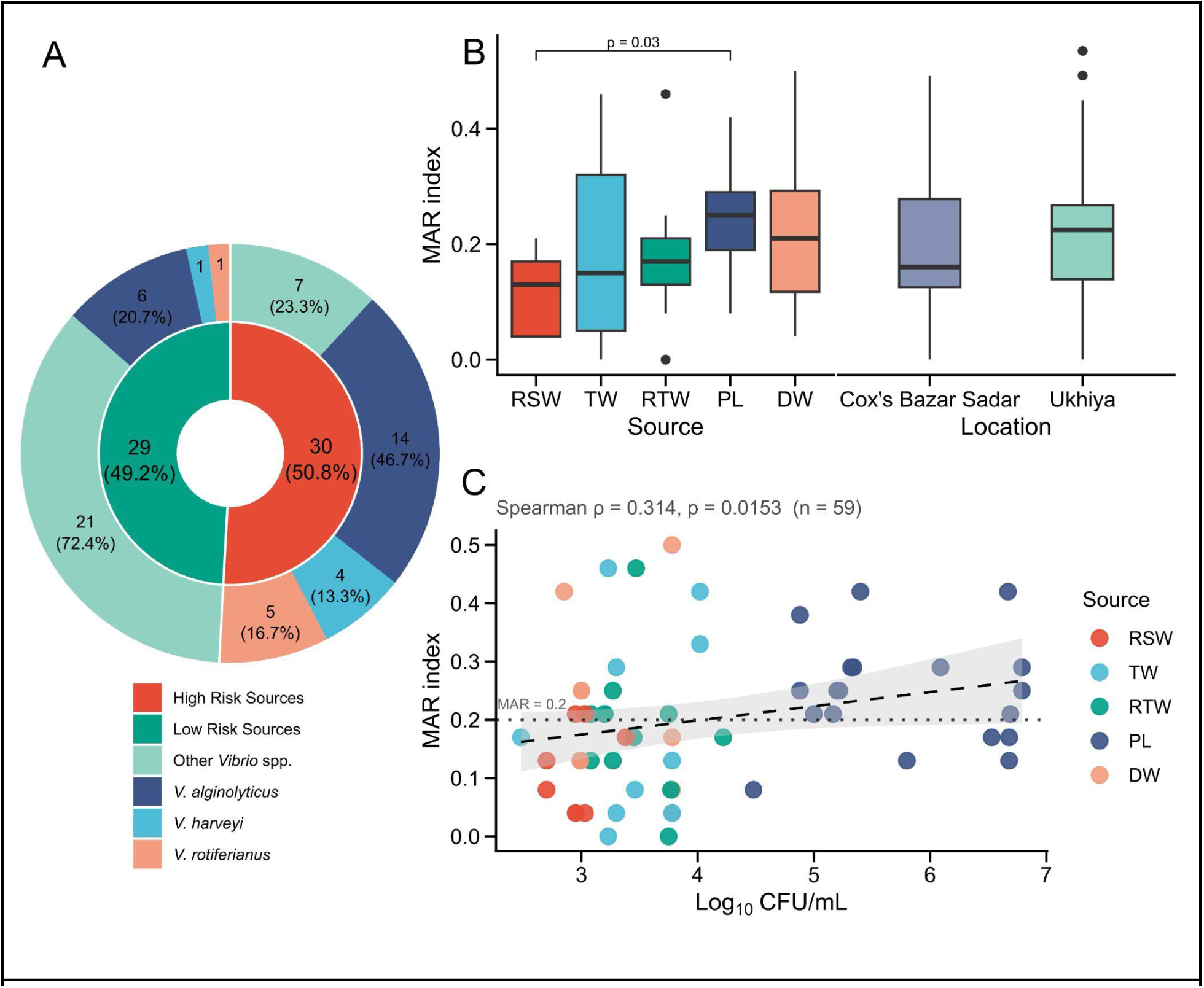
MAR index distribution by species composition, source, location, and its correlation with presumptive *Vibrio* count. (A) Proportion of total isolates (n = 59) by MAR index category, presented in the inner layer as percentages. High risk sources were defined as isolates with MAR index greater than 0.2 and low risk sources were defined with MAR index less than or equal 0.2. The outer layer represents the portion of species composition within each of these categories separately, percentages were calculated relative to each category. (B) Comparative MAR index of *Vibrio* isolates across all sources and locations are presented as a boxplot. Only significant results are indicated here (*P* < 0.05). Statistical significance between sources was calculated using the Kruskal-Wallis test (χ² = 9.64, df = 4, *P* = 0.047), followed by Dunn’s post-hoc pairwise comparisons with *P*-values adjusted using Holm’s method. Only RSW and PL comparison was significant (*P*_adj_ = 0.03). Additionally, Statistical significance between locations was calculated using the Wilcoxon rank sum test (W = 322.5, *P* = 0.7182). (C) Spearman’s rank correlation between the presumptive *Vibrio* count (Log_10_CFU/mL) and MAR index across sources (ρ = 0.314, *P* = 0.0153, n = 59) are presented as a scatter plot. Points are colored by sources.The dashed line represents the linear trend with 95% confidence interval (shaded region). The dotted horizontal line indicates the MAR index threshold (0.2). Abbreviations: DW: discarded water; PL: postlarvae; RSW: raw seawater; RTW: rearing tank water; TW: treated water.

**Table 4.** Multiple antibiotic resistance (MAR) index, resistance patterns, and resistance genes of six most highly resistant *Vibrio* isolates in this study.

| S. no. | Isolate | Source | Location | Phenotypic resistance | Resistant antibiotic class | Resistance genes | MARI |
| --- | --- | --- | --- | --- | --- | --- | --- |
| 01 | *Vibrio alginolyticus* msr8 | DW | Ukhiya | AMP, PRL, CXM, S, TE, CIP, LEV ,OFX , SXT, AZM , E , NA | Penicillins, Cephems, Aminoglycosides, Tetracyclines, Fluoroquinolones, Folate Pathway Inhibitors, Macrolides, Quinolone | *sul2, ermB, tetC* | 0.50 |
| 02 | *Vibrio alginolyticus* msr36 | RTW | Ukhiya | AMP, PRL, CXM, TE , CIP , LEV , OFX, SXT, | Penicillins, Cephems, Tetracyclines, Fluoroquinolones, | *sul2, ermB, tetC* | 0.46 |
|  |  |  |  | AZM, E,<br>NA | Folate Pathway<br>Inhibitors,<br>Macrolides,<br>Quinolone |  |  |
| 03 | *Vibrio*<br>*alginolyticus*<br>msr56 | TW | Cox's<br>Bazar<br>Sadar | AMP, PRL,<br>CXM, TE,<br>CIP, LEV,<br>OFX, SXT,<br>AZM, E,<br>NA | Penicillins,<br>Cephems,<br>Tetracyclines,<br>Fluoroquinolones,<br>Folate Pathway<br>Inhibitors,<br>Macrolides,<br>Quinolone | *sul2, tetC* | 0.46 |
| 04 | *Vibrio*<br>*alginolyticus*<br>msr4 | DW | Ukhiya | AMP,<br>CXM, TE,<br>CIP, LEV,<br>OFX, SXT,<br>AZM, E,<br>NA | Penicillins,<br>Cephems,<br>Tetracyclines,<br>Fluoroquinolones,<br>Folate Pathway<br>Inhibitors,<br>Macrolides,<br>Quinolone | *sul2, ermB, tetC* | 0.42 |
| 05 | *Vibrio* | PL | Ukhiya | AMP , PRL | Penicillins, | – | 0.42 |
|  | *rotiferianus*<br>msr11 |  |  | , AK, S,<br>CIP, LEV,<br>OFX, AZM,<br>E, NA | Aminoglycosides,<br>Fluoroquinolones,<br>Macrolides,<br>Quinolone |  |  |
| 06 | *Vibrio*<br>*alginolyticus*<br>msr6 | PL | Ukhiya | AMP, S,<br>CIP, LEV,<br>OFX, SXT,<br>AZM, E,<br>NA | Penicillins,<br>Aminoglycosides,<br>Fluoroquinolones,<br>Folate Pathway<br>Inhibitors,<br>Macrolides,<br>Quinolone | *sul2, ermB, tetC* | 0.38 |

Abbreviations: Source - DW, discarded water; RTW, rearing tank water; TW, treatment water;
PL:, postlarvae. Antimicrobial agents - AMP, ampicillin; PRL, piperacillin; CXM, cefuroxime sodium; AK, amikacin; S, streptomycin; TE, tetracycline; CIP, ciprofloxacin; LEV, levofloxacin; OFX, ofloxacin; SXT, trimethoprim-sulfamethoxazole; AZM, azithromycin; E, erythromycin; NA, nalidixic acid

The correlation between MAR index and Presumptive *Vibrio* count (Log_10_CFU/mL) were assessed using Spearman’s rank correlation (Figure 7C). The findings indicated a weak to moderate positive relationship between Log_10_CFU/mL and MAR index (ρ = 0.314, *P* = 0.015), suggesting samples carrying higher bacterial loads also tend to carry a greater resistance burden. This pattern was also observed in section 3.1 where PL samples had the highest presumptive *Vibrio* count; consistently, isolates from PL samples in this section also had the highest MAR index.

### 3.7. MDR status of the *Vibrio* isolates

*Vibrio* isolates were categorized as multidrug-resistant (MDR) when an isolate was resistant to at least one antibiotic in three or more antimicrobial classes among 11 antimicrobial classes tested. Among 59 isolates tested, 2 (3.4%) isolates were pan-susceptible, 17 (28.8%) were resistant to one or two classes, 40 (67.8 %) isolates were resistant to 3 or more classes and categorized as MDR. Most of the isolates (n = 14, 23.7%) were resistant to 4 classes and 17 (28.8%) isolates were resistant to 5 or more classes (Figure 8A). Species wise, *V. alginolyticus*, *V. rotiferianus*, and *V. harveyi* accounted for 67.5% (n = 27) of the 40 MDR isolates (Figure 8B). MDR prevalence differed significantly across sample sources (Fisher’s exact test, *P* = 0.016) (Figure 8C). The MDR prevalence increased progressively among the water sources from RSW to DW (44.4% to 62.5%), PL samples had the highest percentage of MDR isolates (94.7%). After pairwise comparison of MDR prevalence among sources, no pairs were found statistically significant (Fisher’s exact test, *P* > 0.05). MDR prevalence also did not differ significantly between the two sampling locations, Cox’s Bazar Sadar and Ukhiya (Fisher’s exact test, *P* = 0.116).

**Figure 8.**
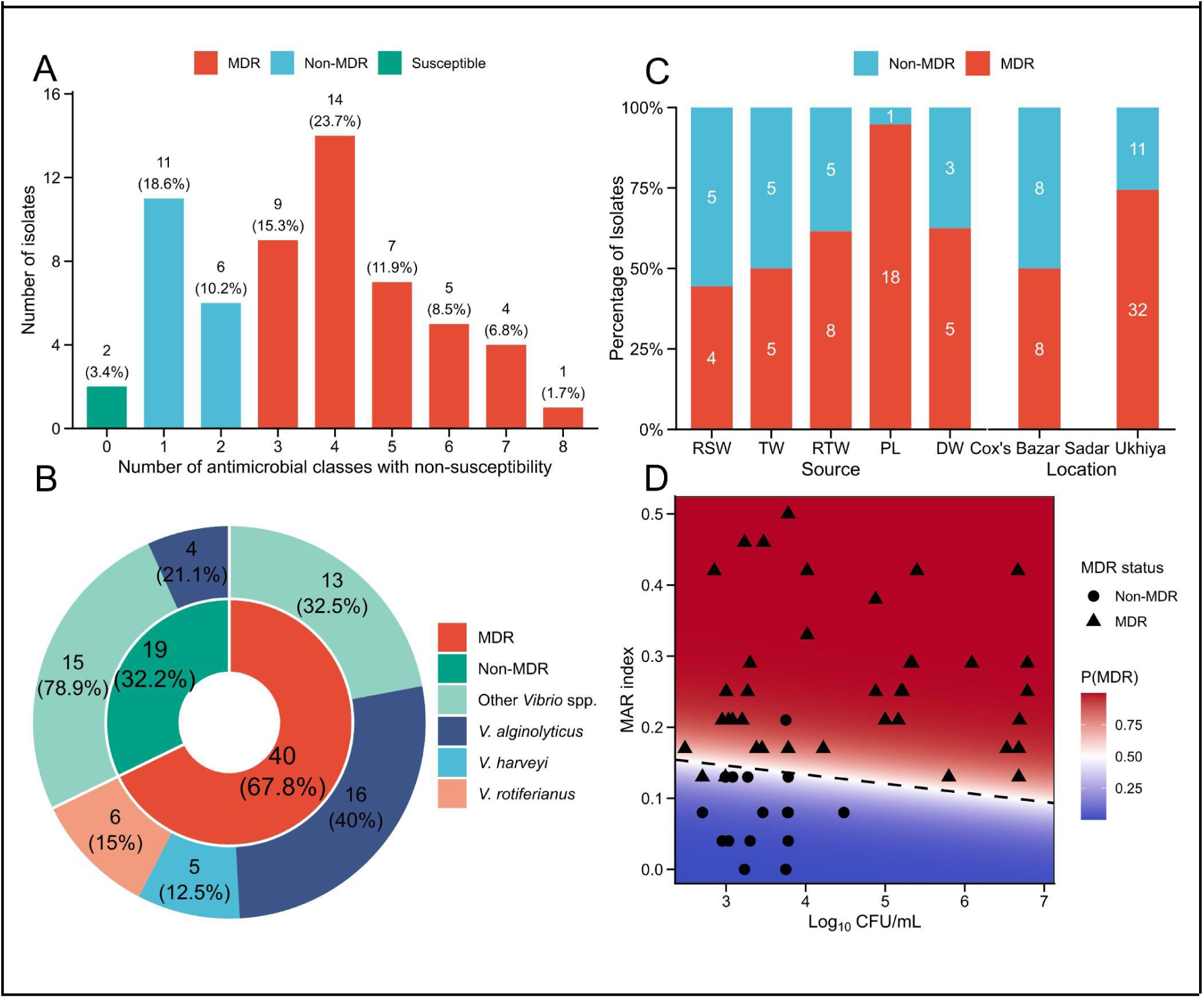
MDR isolate distribution and its relationship with presumptive *Vibrio* count and MAR index. (A) Distribution of isolates by the number of antimicrobial classes with non-susceptibility are presented as a bar plot. An isolate resistant to at least one antibiotic of three or more antimicrobial classes was categorized as MDR. MDR isolates are marked in red, non-susceptible to none are marked as Susceptible (green), and non-susceptible to 1-2 classes are marked as non-MDR (blue). (B) Proportion of total isolates (n = 59) by MDR categories, presented in the inner layer as percentages. The outer layer represents the portion of species composition within each of these categories separately, percentages were calculated relative to each category. (C) Percentages of MDR isolates across sources and locations are presented as bar plots. (D) Logistic regression model (Firth’s penalized likelihood) showing the joint relationship between presumptive *Vibrio* count (Log₁₀CFU/mL), MAR index, and multidrug resistance (MDR) status. The background gradient represents the model-predicted probability of MDR at each combination of CFU and MAR index (blue = low probability, red = high probability). The dashed line indicates the 0.5 decision boundary. Points represent individual isolates (n = 59), shown by their actual observed MDR status (circle = non-MDR, triangle = MDR).

A joint relationship between presumptive *Vibrio* count (Log₁₀ CFU/mL), MAR index, and MDR status was assessed using multiple logistic regression model (Firth’s penalized likelihood) (Figure 8D). The model found the MAR index as a significant predictor of MDR status (β = 39.82, SE = 11.81, *P* < 0.001). In contrast, presumptive *Vibrio* count (Log₁₀CFU/mL) was not found as a significant predictor of MDR status (β = 0.51, SE = 0.41, *P* = 0.191).

### 3.8. Antimicrobial resistance gene prevalence

All the selected *Vibrio* isolates (n = 59) from shrimp hatcheries were screened for five antimicrobial resistance genes, viz., *strA*-*strB*, *sul2*, *ermB*, *tetC*, and *bla*_TEM_ through polymerase chain reaction (PCR). The results were confirmed through agarose gel electrophoresis by detecting correct band size. The prevalence of resistance genes varied significantly among the tested isolates (Chi-square test, χ²=24.264, df = 4, *P* < 0.001). The highest positive result found genes were, *sul2* (20/59, 33.9%), *tetC* (17/59, 28.8%), *strA*-*strB* (9/59, 15.3%), and *ermB* (7/59, 11.9%), respectively (Table 5). In this study, we did not find any positive isolate against *bla*_TEM_. Species containing the highest number of resistant genes were four MDR *V. alginolyticus* isolates (msr4, msr6, msr8, msr36) which were resistant to at least 4 antimicrobial classes, and had 3 resistance genes, viz., *sul2*, *ermB*, *tetC* (Table 4). Among five MDR *V. harveyi* isolates, four of them were positive to *strA*-*strB* and *sul2*. Highest number of *sul2*, *strA*-*strB*, *ermB* genes were found in PL isolates, and RTW isolates had the most number of *tetC* genes, whereas lowest gene prevalence was found in RSW (Supplementary Table 3).

**Table 5.**
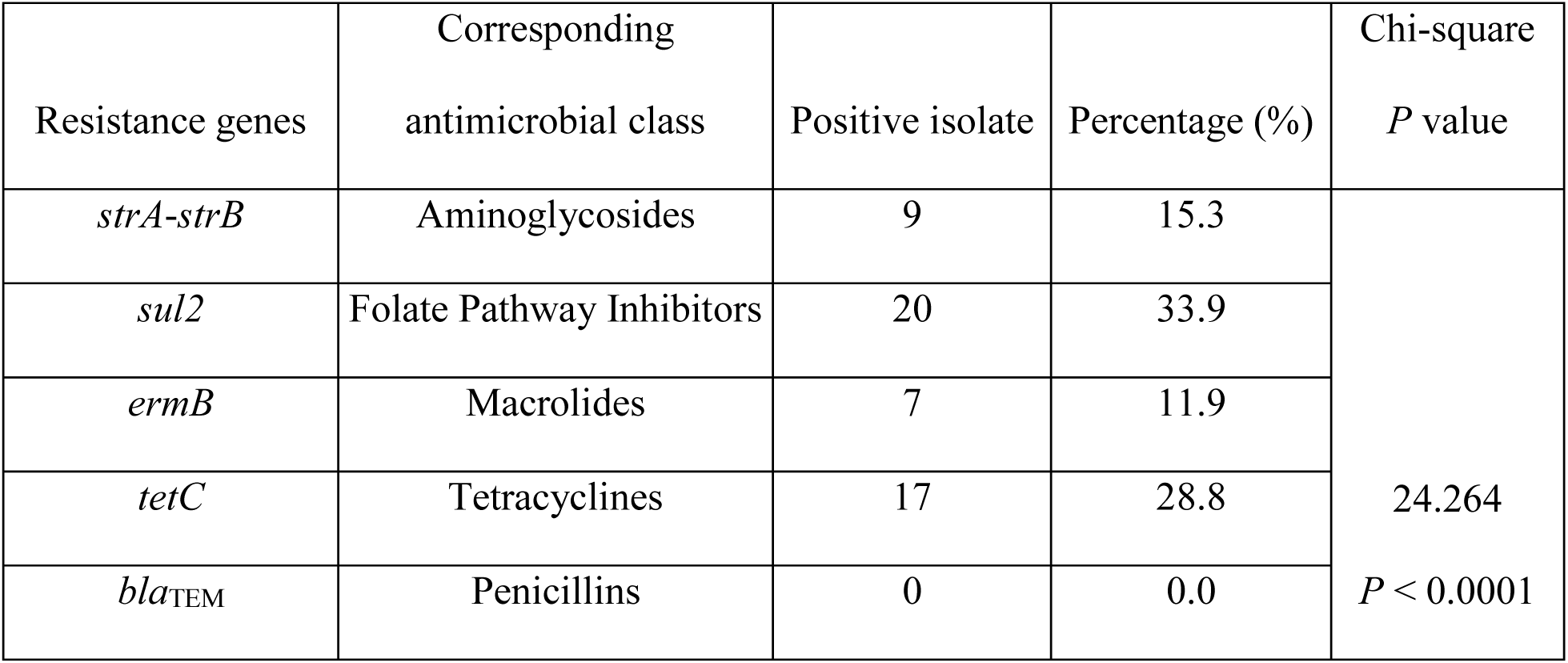
The prevalence of antimicrobial resistance genes (ARGs) in the retrieved *Vibrio* isolates (n = 59) from shrimp hatcheries.

| Resistance genes | Corresponding antimicrobial class | Positive isolate | Percentage (%) | Chi-square<br>*P* value |
| --- | --- | --- | --- | --- |
| *strA-strB* | Aminoglycosides | 9 | 15.3 | 24.264<br>*P* < 0.0001 |
| *sul2* | Folate Pathway Inhibitors | 20 | 33.9 |  |
| *ermB* | Macrolides | 7 | 11.9 |  |
| *tetC* | Tetracyclines | 17 | 28.8 |  |
| *bla*<sub>TEM</sub> | Penicillins | 0 | 0.0 |  |

### 3.9. Association between resistance phenotypes and resistance genes

Due to uneven distribution of isolates across *Vibrio* species, association analysis between phenotypic resistance to different antibiotics could not be performed at species level, hence associations were assessed at the genus level. Associations were assessed by pairwise Fisher’s exact test with Benjamini-Hochberg (BH) correction for multiple comparisons. Three antibiotics (C, CN, F) were excluded from analysis because no isolates were resistant to them. Nineteen pairs of antibiotics showed significant positive associations following Benjamini-Hochberg correction for multiple comparisons (Odds ratio [OR] > 8, including five infinite odds ratios; *P*_adj_ < 0.05) (Table 6). Among these, intra-class associations were found within fluoroquinolones (CIP, OFX, LEV), penicillins (AMP, PRL), and macrolides (AZM, E). Cross-class associations were found between fluoroquinolones (CIP, LEV, OFX) and the quinolone nalidixic acid (NA), the macrolide (AZM), and tetracycline (TE). Cross-class associations were additionally observed between the folate pathway inhibitor trimethoprim-sulfamethoxazole (SXT) and the macrolides (AZM, E), between the quinolone nalidixic acid (NA) and both trimethoprim-sulfamethoxazole (SXT) and tetracycline (TE), and between penicillin (AMP) and cephem (CXM).

**Table 6.** Significant associations between phenotypic resistance to different antibiotics (n = 24) among 59 *Vibrio* isolates determined by Fisher’s exact test with Benjamini-Hochberg (BH) correction for multiple comparisons (*P*_adj_ < 0.05)

| Antibiotics<br>(A<sub>1</sub>-A<sub>2</sub>) | Antimicrobial class | No. of isolates |  |  |  | *P* value | Adjusted<br>*P* value | Odds ratio<br>(OR) | 95% CI |
| --- | --- | --- | --- | --- | --- | --- | --- | --- | --- |
|  |  | S-S | S-R | R-S | R-R |  |  |  |  |
| CIP-OFX | Fluoroquinolones | 41 | 2 | 3 | 13 | 1.28E-08 | 2.69E-06 | 74.49 | 10.82-977.66 |
| LEV-OFX | Fluoroquinolones | 44 | 6 | 0 | 9 | 3.98E-07 | 4.18E-05 | Inf | 10.34-Inf |
| CIP-LEV | Fluoroquinolones | 43 | 0 | 7 | 9 | 9.10E-07 | 6.37E-05 | Inf | 8.98-Inf |
| AMP-CXM | Penicillins-Cephems | 21 | 1 | 12 | 25 | 1.46E-06 | 7.69E-05 | 40.88 | 5.38-1858.51 |
| LEV-NA | Fluoroquinolones-Quinolones | 41 | 9 | 0 | 9 | 3.87E-06 | 1.63E-04 | Inf | 7.01-Inf |
| OFX-NA | Fluoroquinolones-Quinolones | 38 | 6 | 3 | 12 | 5.14E-06 | 1.80E-04 | 23.22 | 4.61-167.46 |
| LEV-AZM | Fluoroquinolones-Macrolides | 39 | 11 | 0 | 9 | 1.34E-05 | 4.01E-04 | Inf | 5.63-Inf |
| CIP-NA | Fluoroquinolones-Quinolones | 37 | 6 | 4 | 12 | 1.80E-05 | 4.72E-04 | 17.17 | 3.75-100.28 |
| AMP-PRL | Penicillins | 21 | 1 | 17 | 20 | 7.65E-05 | 0.0018 | 23.55 | 3.14-1065.33 |
| CIP-AZM | Fluoroquinolones-Macrolides | 35 | 8 | 4 | 12 | 1.01E-04 | 0.0021 | 12.37 | 2.85-67.65 |
| TE-CIP | Tetracyclines-Fluoroquinolones | 41 | 8 | 2 | 8 | 1.93E-04 | 0.0036 | 19.00 | 3.06-215.46 |
| SXT-E | Folate Pathway<br>Inhibitors-Macrolides | 20 | 22 | 0 | 17 | 2.08E-04 | 0.0036 | Inf | 3.13-Inf |
| TE-LEV | Tetracyclines-Fluoroquinolones | 46 | 3 | 4 | 6 | 3.19E-04 | 0.0052 | 20.74 | 3.14-184.30 |
| OFX-AZM | Fluoroquinolones-Macrolides | 35 | 9 | 4 | 11 | 3.77E-04 | 0.0056 | 10.15 | 2.34-54.74 |
| SXT-AZM | Folate Pathway<br>Inhibitors-Macrolides | 34 | 8 | 5 | 12 | 4.70E-04 | 0.0066 | 9.69 | 2.38-46.48 |
| TE-NA | Tetracyclines- | 39 | 10 | 2 | 8 | 6.04E-04 | 0.0075 | 14.67 | 2.43-162.63 |
|  | Quinolones |  |  |  |  |  |  |  |  |
| SXT-NA | Folate Pathway<br>Inhibitors-Quinolones | 35 | 7 | 6 | 11 | 5.69E-04 | 0.0075 | 8.72 | 2.15-40.38 |
| AZM-E | Macrolides | 19 | 20 | 1 | 19 | 9.93E-04 | 0.0116 | 17.34 | 2.30-785.33 |
| TE-OFX | Tetracyclines-Fluoroquinolones | 41 | 8 | 3 | 7 | 0.0015 | 0.0161 | 11.25 | 2.06-82.27 |

Note: S = susceptible, R = resistant; column order is A<sub>1</sub>–A<sub>2</sub>.
Abbreviations: AMP, ampicillin; AMC, amoxicillin-clavulanic acid; NA, nalidixic acid; PRL, piperacillin; TZP, piperacillin-tazobactam; CXM, cefuroxime sodium; FEP, cefepime; CTX, cefotaxime; FOX, cefoxitin; CAZ, ceftazidime; CRO, ceftriaxone; IPM, imipenem; AK, amikacin; S, streptomycin; TE, tetracycline; CIP, ciprofloxacin; LEV, levofloxacin; OFX, ofloxacin; SXT, trimethoprim-sulfamethoxazole; AZM, azithromycin; E, erythromycin

Each resistance gene was tested against its biologically expected phenotypes: *sul2* with trimethoprim-sulfamethoxazole (SXT), *ermB* with erythromycin (E) and azithromycin (AZM), *tetC* with tetracycline (TE), and *strA*-*strB* with streptomycin (S) (Table 7). As there was no positive isolate against *bla*_TEM_, so it was excluded from the analysis. A strong, statistically significant positive association was found between SXT resistance and the presence of the *sul2* gene (OR = Inf, 95% CI: 29.87-Inf; *P*_adj_ = 2.05E-11), and a moderate positive significant association was found between streptomycin (S) resistance and the presence of the *strA*-*strB* gene (OR = Inf, 95% CI: 1.86-Inf; *P*_adj_ = 0.0083). Azithromycin (AZM) resistance and presence of *ermB* gene showed positive associations (OR = 5.95, *P* = 0.0383), which did not remain significant following Benjamini-Hochberg (BH) correction for multiple comparisons (*P*_adj_ = 0.0639). Positive associations were also found between erythromycin (E) and *ermB* (OR = 3.40), and between tetracycline (TE) and *tetC* (OR = 3.02), but the associations were not statistically significant (*P*_adj_ > 0.05).

**Table 7.** Associations among antimicrobial resistance genes (n = 4) and its corresponding antibiotic determined by Fisher’s exact test with Benjamini-Hochberg (BH) correction for multiple comparisons.

| Resistance gene | Corresponding antibiotic | No. of isolates |  |  |  | *P* value | Adjusted *P* value | Odds ratio (OR) | 95% CI |
| --- | --- | --- | --- | --- | --- | --- | --- | --- | --- |
|  |  | G(-)_A(S) | G(-)_A(R) | G(+)_A(S) | G(+)_A(R) |  |  |  |  |
| *sul2* | SXT | 39 | 0 | 3 | 17 | 4.11E-12 | 2.05E-11 | Inf | 29.87-Inf |
| *strA-strB* | S | 26 | 24 | 0 | 9 | 0.0033 | 0.0083 | Inf | 1.86-Inf |
| *ermB* | AZM | 37 | 15 | 2 | 5 | 0.0383 | 0.0639 | 5.95 | 0.86-69.14 |
| *tetC* | TE | 37 | 5 | 12 | 5 | 0.1332 | 0.1664 | 3.02 | 0.59-15.72 |
| *ermB* | E | 19 | 33 | 1 | 6 | 0.4045 | 0.4045 | 3.40 | 0.37-166.99 |

Note: G(+)/G(-), presence/absence of resistance gene; A(S)/A(R), susceptible/resistance of the corresponding antibiotics.
Abbreviations: SXT, trimethoprim-sulfamethoxazole; S, streptomycin; AZM, azithromycin; TE, tetracycline; E, erythromycin

When association analyses were performed among resistance genes, *strA*-*strB* and *sul2* showed a significant positive association (OR = 23.7, *P*_adj_ = 0.002), *sul2* and *ermB* also showed a significant positive association (OR = 15.46, *P*_adj_ = 0.014) (Supplementary Table 4). The *sul2*-*tetC* gene pair (OR = 3.10) and *ermB*-*tetC* gene pair (OR = 3.89) were positively associated but not significantly (*P*_adj_ > 0.05). The remaining gene pairs, *strA*-*strB* and *tetC*, *strA*-*strB* and *ermB* were negatively associated and produced non-significant results (OR = 0, *P*_adj_ > 0.05).

## 4. Discussion

Vibrios are broadly distributed across aquatic habitats spanning from freshwater and estuarine to marine waters and widely prevalent in the aquaculture operations worldwide (de Souza Valente & Wan, 2021). Some are part of the normal flora, while others are capable of causing vibriosis in aquaculture systems and pose zoonotic risks to humans (Austin, 2010). Emerging disease outbreaks and antibiotic resistance mechanisms of *Vibrio* species have made treatment increasingly difficult. Their ubiquitous presence and transmission of resistance genes through horizontal gene transfer (HGT) have imposed a threat to the One Health approach (Marques et al., 2022). These characteristics have gained the attention of researchers to study their species composition within a given environment and antibiotic resistance patterns. Among numerous *Vibrio* studies conducted across different aquaculture settings, studies specifically focused on hatchery systems are rare. Studies on *Vibrio* species composition and antibiotic resistance from hatchery systems can help track and prevent vibriosis outbreaks, understand *Vibrio* ecology, and current status of antibiotic resistance and its potential transmission to the environment, other aquaculture settings, and public health. In the present study, we collected samples, viz., raw seawater, treated water, rearing tank water, discarded water, and postlarvae from the shrimp hatcheries of Cox’s Bazar, Bangladesh. The collected samples were analyzed to identify *Vibrio* species and their phenotypic resistance and resistance genes along the hatchery water flow system and postlarvae. This study may contribute useful information from Bangladeshi shrimp hatcheries toward understanding broader marine *Vibrio* diversity and the current status of antibiotic resistance in this setting.

The presumptive *Vibrio* species count from TCBS plates of each sample revealed a significantly higher count in the PL samples than the water samples (Figure 2). This is consistent with reports that the shrimp postlarvae harbors more *Vibrio* species than the surrounding water, driven by host selectivity (Garibay-Valdez et al., 2020). This might also be due to more favorable conditions provided by the host, including nutrients and surfaces for attachment and biofilm formations than the surrounding waters providing unfavorable conditions for bacterial proliferation (de Souza Valente & Wan, 2021). TCBS media is highly selective for *Vibrio* species isolation due to its high alkaline pH and bile salts, colony colors on this media (green and yellow) are based on sucrose-fermenting capability of the species (Kaysner et al., 2004). The observed discordance between morphological traits of the *Vibrio* isolates on the TCBS plates and species identification is due to the limited discriminatory power of this medium. Again, some colonies showed standalone characteristics in this study including *V. alginolyticus* colonies (Figure 3), which might help in predicting probable species rather than confirming them. This also underscores the limitation of dependence on morphological traits alone to identify *Vibrio* species. Consequently, 16S rRNA gene sequencing was used to identify the bacterial isolates from the shrimp hatcheries. All the bacterial isolates were confirmed to belong to the *Vibrio* genus, comprising 19 different species belonging to nine clades (Figure 4). Among them, the Harveyi clade species (38/59) were the most dominant, viz., *V. alginolyticus* (20/59), *V. rotiferianus* (6/59), *V. harveyi* (5/59), *V. natriegens* (3/59), *V. campbellii* (2/59), and *V. parahaemolyticus* (2/59). The maximum-likelihood phylogenetic tree placed all the identified isolates close to their type strain, and the isolates from the same clade were placed close to each other. The ultrafast bootstrap and SH-aLRT values were low for the nodes separating the clade members. The branch length among the clade species was minimal, reflecting lower diversity at the 16S locus. This is consistent with the high sequence similarity (>97%) known to limit the species-level identification of *Vibrio* species based on 16S rRNA gene alone (Sawabe et al., 2007). *V. alginolyticus* was the most identified species and the only species that was present in the five sampling sources from raw seawater through postlarvae to discard water, suggesting it is well adapted to persist throughout the hatchery system (Figure 5). *V. alginolyticus* was also identified as the most dominant species from shrimp breeding grounds of China (Yu et al., 2023). *V. alginolyticus* is a common marine bacterium; it is also responsible for various shrimp postlarvae diseases such as zoea II syndrome and septic hepatopancreatic necrosis disease (de Souza Valente & Wan, 2021). *V. harveyi* and *V. campbellii* were also recovered from the hatchery system. These two species are among the major causes of luminous vibriosis disease in shrimp hatchery (Kumar et al., 2021; Lavilla-Pitogo et al., 1990). Most of the *V. harveyi* isolates (4/5) were identified from shrimp PL, suggesting a potential risk of luminous vibriosis outbreak at this life stage. *V. rotiferianus* was also identified in substantial numbers in this study. It has also been reported to cause diseases in shrimps (Haifa-Haryani et al., 2023; Zhang et al., 2024). *V. tubiashii*, *V. natriegens*, and *V. proteolyticus* were each identified in three isolates. *V. tubiashii* has been reported to be involved with disease outbreak in shellfish hatchery (Hasegawa & Häse, 2009). Although it has not been reported to cause any shrimp diseases specifically, further investigations are required about its role in the shrimp hatchery system. *V. natriegens* is a fast-growing marine bacterium that can produce a biosurfactant capable of reducing pathogenicity of the shrimp pathogen *V. harveyi* (Kannan et al., 2019). *V. proteolyticus* has the ability to infect banana prawn (*Fenneropenaeus merguiensis*) under experimental conditions (Zhong et al., 2023). *V. parahaemolyticus* was found in small numbers (2/59), but it is widely recognized as a shrimp larval pathogen (de Souza Valente & Wan, 2021). The remaining *Vibrio* species were each identified in one or two isolates, reflecting the broader marine *Vibrio* diversity in the shrimp hatchery system. Apart from being shrimp pathogens, some vibrios are also recognized as zoonotic pathogens capable of infecting humans (Austin, 2010). In the present study, four species including *V. parahaemolyticus*, *V. harveyi*, *V. alginolyticus*, and *V. fluvialis* have zoonotic potential. *V. parahaemolyticus* is reported to cause gastroenteritis, wound infection, and sepsis in humans (Rezny & Evans, 2023). *V. fluvialis* is reported to cause vomiting and diarrhoea, frequently with bloody stools (Forsythe et al., 2009). *V. harveyi* can infect humans through wounds and cause severe inflammatory responses (Yu et al., 2022). *V. alginolyticus* can cause wound or ear infections (Giannella, 2010). Zoonotic *Vibrio* species can transmit through seafood consumption, direct seawater contact, or handling of infected animals (Baker-Austin et al., 2018), making shrimp hatcheries as a potential medium for zoonotic disease transmission. Overall this diverse *Vibrio* composition reflects marine ecological complexity, with potential risks to the disease outbreaks in the postlarvae and zoonotic risk to public health. Given their potential disease risk, antimicrobial resistance screening is essential for guiding treatment and biosecurity strategies in shrimp hatchery systems.

The extensive use of antibacterials has led to the emergence of antimicrobial-resistant bacteria in the aquaculture systems, which poses risks to the public health, veterinary medicine, and environmental safety (Odoi et al., 2025). Consequently, frequent assessment of antimicrobial resistance in the aquaculture systems is required to enable in-depth analysis of possible threats to both animal and public health (Milijasevic et al., 2024). In the present study, fifty nine *Vibrio* isolates were tested against 24 antibiotics from eleven antimicrobial classes selected from both human clinical and aquaculture perspectives. The highest resistance phenotype was observed for erythromycin (E) (66.1%), ampicillin (AMP) (62.7%), and streptomycin (S) (55.9%) (Figure 6B). Erythromycin inhibits further growth of bacteria by inhibiting protein synthesis, whereas ampicillin kills bacteria by disrupting cell wall synthesis (Farzam et al., 2023; Sabunwala et al., 2026). Erythromycin and ampicillin are among the most frequently used antibiotics in shrimp hatcheries and farms in Bangladesh, their frequent usage might have led to the observed antimicrobial resistance (Bashar et al., 2026; Kawsar et al., 2026). Macrolides (azithromycin and erythromycin) are highly effective against Gram-positive bacteria and they have limited activity against Gram-negative bacteria (Wangkahart et al., 2025). Nevertheless, azithromycin showed higher sensitivity and erythromycin showed lower sensitivity in this study. Azithromycin is a 15-membered-ring macrolide which showed better antimicrobial activity against gram-negative bacteria than the 14-membered-ring macrolide erythromycin (Retsema et al., 1987). Overall, erythromycin’s high resistance may be attributable partly to these structural differences and partly to higher selection pressure. Similar pattern of high erythromycin resistance and comparatively moderate ampicillin resistance were reported in diverse bacteria from shrimp PL nurseries in Bangladesh (Yasin et al., 2022) and in *Vibrio* species recovered from selected freshwaters in Southwest Nigeria (Adesiyan et al., 2022). Streptomycin does not appear among the antibiotics that are used in Bangladeshi shrimp hatchery and farm (Bashar et al., 2026; Kawsar et al., 2026). Although streptomycin is not reported to be used, its high resistance may reflect legacy selection pressure as it is one of the earliest antibiotics and its resistance recorded in the late 1940s (Afroze et al., 2025). As resistant genes often persist after withdrawal, due to minimal fitness cost to the host bacterium, historical exposure may contribute to the current streptomycin resistance (Tamminen et al., 2011). Similar streptomycin resistance was also reported by other authors (He et al., 2016; Kathleen et al., 2016).

In the present study, no isolates were found resistant to chloramphenicol (C), gentamicin (CN), and nitrofurantoin (F) (Figure 6B). Chloramphenicol obstructs the synthesis of bacterial protein by attaching to the 50S ribosomal subunit; gentamicin binds to the 16S rRNA at the 30S ribosomal subunit, disturbs the mRNA translation; and nitrofurantoin is reduced by bacterial flavoproteins into reactive intermediates that disrupt DNA, RNA, and cell wall synthesis (Beganovic et al., 2018; Giedraitiene et al., 2022; Syroegin et al., 2022). Chloramphenicol and nitrofurantoin exhibit strong antibacterial activity against both Gram-positive and Gram-negative bacteria, whereas gentamicin is primarily used against Gram-negative bacteria (Bradley & Sauberan, 2012; Moore et al., 1984; Papich, 2013). Bangladesh faced an export crisis during 2009 due to contamination of banned antibiotics nitrofuran (including nitrofurantoin) and chloramphenicol in shrimps and prawns. Consequently, various reforms were adopted in this sector by the government. This led to the decreasing trend of the presence of nitrofuran and chloramphenicol in shrimps (Hassan et al., 2013). Although limited use of chloramphenicol has been reported, it is not one of the routinely used antibiotics in Bangladesh shrimp hatcheries (Kawsar, 2026). This declining and limited usage supports the idea that lower selection pressure leads to lower antibiotic resistance (Oz et al., 2014; Zhang et al., 2015). Higher susceptibility to chloramphenicol and gentamicin was also observed in other studies (Changsen et al., 2023; Kathleen et al., 2016; Rahman et al., 2020), while higher susceptibility to nitrofurantoin was reported by Rahman et al. (2020).

In the present study, variation coefficient percentage (VC%) was calculated to understand the variability of the antibiotic activity against the *Vibrio* isolates (Figure 6A). The higher VC% indicates that the antibiotics has a variable activity against the tested *Vibrio* isolates, therefore it’s in vivo antimicrobial activity, whether resistant or susceptible is more unpredictable. On the other hand, the lower VC% indicates that the antibiotic has more stable and uniform activity against the tested *Vibrio* isolates, therefore its in-vivo antimicrobial activity, whether resistant or susceptible is more predictable (Lakhssassi et al., 2005). Higher VC% (>25%) was observed for 12 antibiotics, viz., AMP (125.8%), E (70%), SXT (66.2%), S (63%), NA (61%), AZM (57.7%), OFX (46.2%), CIP (44.6%), LEV (43.6%), CXM (39.4%), PRL (37.1%), and TE (34.6%), meaning these antibiotics activity is uncertain against the hatchery *Vibrio* isolates. Ampicillin (AMP) had the highest VC% of 125.8%, which is considerably higher than the VC% for ampicillin reported by Sony et al. (2021), where TE and SXT also showed high variability (>25%). Older antibiotics such as ampicillin, erythromycin, streptomycin, nalidixic acid, and tetracycline showed higher VC% in this study, which is consistent with the findings of (Lakhssassi et al., 2005; Sony et al., 2021) that older drugs are associated with higher VC%. But several newer generation antibiotics such as ciprofloxacin, levofloxacin, ofloxacin, and azithromycin also showed higher VC%, indicating antibiotic age alone may not fully explain variability in activity against *Vibrio* isolates from this hatchery setting. Final activity score was assigned based on both susceptibility percentage (S%) and variation coefficient percentage (VC%), ranging from 0 to 8, which was classified as inactive (0), poorly active (≥1 to <5), fairly active (≥5 to <8), or very active (8) (Sony et al., 2021). In this study, none of the 24 tested antibiotics were categorized as inactive (0) or very active (8) (Figure 6D). They were categorized as fairly active or poorly active. Seven antibiotics, viz., CXM, NA, SXT, AZM, E, S, and AMP were categorized as poorly active, suggesting that treating vibriosis with these antibiotics in these shrimp hatcheries may not be successful. Again, five antibiotics (TE, PRL, CIP, LEV, and OFX) had high VC%, yet they did not fall under the poorly active category, because their high susceptibility percentage compensated for high VC%, illustrating why VC% alone should not be used to judge an antibiotic’s overall reliability.

The multiple antibiotic resistance (MAR) index of 59 *Vibrio* isolates was calculated as described by Krumperman (1983), where a value greater than 0.2 is an indication that the isolate is likely from a high antibiotic selection pressure environment, and a value less than or equal to 0.2 is an indication that the isolate is likely from a low antibiotic selection pressure environment. In the present study, the MAR index ranged from 0 to 0.5. Of the 59 isolates, 30 (50.8%) had a MAR index greater than 0.2 with a median of 0.21. MAR index of *Vibrio* isolates from water samples showed an increasing trend from RSW to DW, and PL isolates showed the highest value, with RSW and PL showing a significant difference (Figure 7B). Notably, the proportion of RSW isolates with a MAR index greater than 0.2 was 22.2% (2/9), which is comparatively lower than the levels reported for other geographical coastal waters, including Saudi Arabia, Malaysia, and Tunisia (Elhadi et al., 2022; You et al., 2016; Zaafrane et al., 2022), suggesting that incoming seawater was not a heavily contaminated source of antibiotic-resistant *Vibrio*. The lowest observed MAR values of RSW, along with higher MAR values of TW, RTW, PL, and DW suggests that the hatchery systems are likely under high antibiotic selection pressure. In hatchery systems, preventative antibiotic therapy is more common than in grow-out phases, and it is documented that Bangladeshi shrimp hatcheries use up to 80 kg of antibiotics per rearing cycle (Hinchliffe et al., 2018; Thornber et al., 2020). Among the water sources (RSW, TW, RTW, DW), DW had the highest MAR index, suggesting that antibiotic-resistant *Vibrio* originating within the hatchery systems accumulate in discharge water. This aligns with reports that Bangladeshi aquaculture infrastructure, including shrimp hatcheries, lacks adequate effluent treatment systems, resulting in direct disposal of hatchery wastes into the seawater with minimal or no effluent treatment (Haque & Mahmud, 2025). This poses a serious threat to marine environment through disposing of hatchery wastes containing antibiotic resistant *Vibrio*. This practice may contribute to the wider spreading of resistant strains in the marine environment, posing a potential risk to both marine ecosystems and public health through direct water contact and seafood consumption. Among studies conducted in Bangladeshi shrimp aquaculture systems, the proportion of isolates exceeding MAR index threshold of this study (50.8%) is higher than the findings of Rahman et al. (2020), who found 18.75% of *Vibrio* spp. from shrimp farms exceeding this threshold. On the other hand, it is similar to Sohidullah et al. (2025) and Haque et al. (2023), who found 51.28% of *V. parahaemolyticus* and 53.9% of *Vibrio* spp., respectively, from shrimp farms exceeding this threshold. Furthermore, it is lower than the findings of Yasin et al. (2022), who found 86.7% of diverse bacteria from shrimp PL nurseries exceeding this threshold. Among studies conducted in other Asian shrimp aquaculture systems, the proportion of isolates exceeding the MAR threshold in this study is lower than that reported by Letchumanan et al. (2015) in Malaysia (83%), Narsale et al. (2024) in India (68%), Silvester et al. (2015) in India (72%), and Yu et al. (2023) in China (63.5%). These variations in MAR values of the *Vibrio* isolates might be due to differences in antibiotic selection pressure from different sources and geographical locations (Lesley et al., 2011; Tunung et al., 2012). Both the presumptive *Vibrio* count and the MAR index had the highest values in the PL samples. Therefore, their relationship was evaluated using Spearman’s rank correlation analysis (Figure 7C). The results revealed a weak-to-moderate significant correlation between them (Spearman correlation, ρ = 0.314, *P* = 0.0153, n = 59), indicating that samples with high *Vibrio* loads also tend to have higher MAR index. This finding contradicts the findings of Crettels et al. (2023), who found no significant relationship between bacterial abundance and the MAR index in freshwater *E. coli* from Belgian bathing waters (Pearson correlation, *P* = 0.56, n = 54). This discrepancy might be due to species diversity and environmental settings. Additionally, the sample types were also different, they sampled only water, whereas our study incorporated both water and PL samples, which may include host associated resistance dynamics not present in their water only samples. Further study is required to clarify the relationship between bacterial abundance and MAR index across different aquatic and aquaculture settings.

The antibiotics resistance pattern abundance (ARPA) is the ratio of total resistance types to total studied bacterial isolates, which was originally given by Deng et al. (2020). Several other authors have adapted this calculation to their studies (Adesiyan et al., 2021; Adesiyan et al., 2022; Ferri et al., 2024; Ferri et al., 2025; S. J. et al., 2026; Yu et al., 2023). The reported ARPA values in the literature ranged from 0.106 to 0.714. In the present study, the ARPA value of *Vibrio* isolates (n = 59) from shrimp hatcheries of Bangladesh was 0.78 overall and 0.8 for *V. alginolyticus* (n = 20) specifically (Table 4), which are the highest reported up to this date. These varying resistance patterns from different studies suggest bacteria exhibit different resistance patterns based on antibiotic selection pressure of each environment (Adesiyan et al., 2022; Deng et al., 2020). The high ARPA values from shrimp hatchery also suggests these resistance patterns are likely to be originating from multiple sources within the hatchery system, which will make it difficult to manage through a single type of intervention.

Environment, aquaculture, and public health are under constant threat by globally rising multi-drug resistant (MDR) bacterial strains. Regular antibiotic sensitivity testing is essential for the screening of increasing MDR isolates to choose the correct antibiotic for disease control (Afroze et al., 2025). In the present study, an isolate was considered MDR, when it was found resistant to three or more antimicrobial classes. A total of 40 of 59 (67.8%) *Vibrio* isolates were categorized as MDR (Figure 8B). This is higher than the previously reported *Vibrio* MDR prevalence from Bangladeshi shrimp aquaculture and retail market (Haque et al., 2023; Sohidullah et al., 2025; Sultana et al., 2025). This discrepancy might be due to different rearing condition and antibiotics selection pressure between the hatcheries and farms; further studies are required to confirm this. When compared with other global study for *Vibrio* spp. from marine environments, our MDR result was higher than several studies (Banerjee & Farber, 2018; Ferri et al., 2024; Jeamsripong et al., 2020; Sony et al., 2021; Zhao et al., 2018). And it was lower than several studies (Algammal et al., 2025; Elhadi et al., 2022; S. J. et al., 2026). Compared with other Bangladeshi and global studies, MDR prevalence of this study falls in the upper-middle range. Bangladesh lacks a national antibiotic usage policy for aquaculture (Luthman et al., 2024), and without proper regulation of antibiotics usage, this prevalence might rise over time. This elevated MDR prevalence might threaten the animal, environmental, and public health seriously. At species level, *V. alginolyticus*, *V. rotiferianus*, and *V. harveyi* comprised 52.5% (31/59) of the total *Vibrio* isolates, yet they accounted for 67.5% (27/40) of MDR isolates (Figure 7A) and 76.7% (23/30) of isolates exceeding MAR threshold (Figure 8B), suggesting these species are most likely carrying the antibiotic resistance burden in the hatchery environment. In the present study, no isolate was categorized as extensive drug resistant (XDR), defined as resistant to all but ≤2 antimicrobial classes (Magiorakos et al., 2012). Four isolates were resistant to 7 antimicrobial classes, and one isolate was resistant to 8 antimicrobial classes (Figure 8A), all of these isolates were identified as *V. alginolyticus* and they were short one to two classes to meet the criteria of XDR. If proper antibiotic management strategies are not adopted, these isolates might turn into extensive drug resistant (XDR) or even pan-drug resistant (PDR). Among the MDR prevalence across different hatchery sources, the PL source showed the highest MDR prevalence at 94.7% (18/19) (Figure 8C), and the MAR index also showed the highest for PL source compared to other sources (Figure 7B). Both PL and RTW originated from the same tank, yet PL has a higher MAR index and MDR prevalence than RTW. This pattern may be explained through several factors. During the larval rearing period, water is exchanged frequently about 30-40% per day (Kungvankij et al., 1985), so the microbial load is getting diluted. On the other hand, PL are highly prone to disease from the pathogenic bacterial strains and treated with antibiotics (Sotomayor et al., 2019), resulting in longer exposure to the antibiotics. Artemia is fed to the PL as live feed and is known to carry high *Vibrio* density (Vaseeharan & Ramasamy, 2003), which may also introduce antibiotic resistant *Vibrio* to the PL. PL may provide favorable nutrition and shelter for bacterial attachment and biofilm formation (Ashrafudoulla et al., 2019). *Vibrio* species can switch between free-living (planktonic) and biofilm lifestyles, and biofilm formation can increase antibiotic resistance compared to free-living state (Meza-Villezcas et al., 2022). These factors may explain the elevated antibiotic resistance observed in the PL compared to RTW. The post-larvae (PL) produced in Cox’s Bazar hatchery are transported to the southern part of the country including Bagerhat, Khulna, and Satkhira districts for the grow-out phases (Debnath et al., 2016). In these regions, traditional gher system is common, where ghers exchange water from tidal channel, river, estuary, and directly from ocean with little or no water treatment (Thornber et al., 2026). When these PL, carrying a high prevalence of MDR *Vibrio*, are released to grow-out farms, they may also introduce antibiotic-resistant strains into the farm environment. As these traditional ghers lack biosecurity measures, these MDR *Vibrio* isolates may spread through the wild and cultured animals, the riverine environment, and the human food chain, posing a threat under a One Health framework.

In the present study, presumptive *Vibrio* count, MAR index, and MDR prevalence were all highest in PL samples compared to the water sources, suggesting a possible relationship between higher presumptive *Vibrio* loads, MAR index, and MDR prevalence. Additionally, in the 3.5 section a weak-to-moderate significant relationship was found between presumptive *Vibrio* loads and MAR index. Given this, the joint relationship between presumptive *Vibrio* count, MAR index, and MDR status was assessed using a multivariable logistic regression model with Firth’s penalized likelihood method (Figure 8D). The model identified MAR index as a significant predictor of MDR status. In contrast, presumptive *Vibrio* count was not found as a significant predictor of MDR status. Although there was a weak correlation between presumptive *Vibrio* loads and the MAR index, resistance burden became the sole predictor of MDR status rather than bacterial load. Statistical analyses of MDR data have generally relied on descriptive approaches rather than regression-based modeling (Agga & Scott, 2015), and to our knowledge, no previous study has evaluated the joint relationship between these variables using a multivariable logistic regression model in *Vibrio* isolates from aquaculture settings. MAR index has been previously used to predict resistance phenotypes in other bacterial species and clinical settings using this kind of modeling (Yildiz et al., 2025), but not in combination with bacterial load, or in relation to MDR status. This underscores the value of the present modeling approach for future AMR surveillance studies in aquaculture systems.

*Vibrio* species harboring antibiotic-resistant genes (ARGs) represent an alarming trend, as they can easily disseminate resistance genes both within their own genus (Deng et al., 2019) and to other genera (Nonaka et al., 2022). This poses a risk at the hatchery stage with potential to propagate through PL transfer to grow-out farms and the human food chain. In the present study, five antibiotic resistance genes including *sul2*, *ermB*, *tetC*, *strA*-*strB*, and *bla*_TEM_ were screened using polymerase chain reaction (PCR). Unlike other intrinsic resistance genes that are integrated into the *Vibrio* chromosome, these genes are acquired through horizontal gene transfer (HGT) via extrachromosomal replicons, integrons, transposable elements, and SXT/R391 integrative conjugative elements (De, 2021; Furushita et al., 2003; Urban-Chmiel et al., 2022). The *sul2* gene encodes an alternative dihydropteroate synthase (DHPS) enzyme that is insensitive to sulfonamides, allowing the bacteria to continue folate synthesis in the presence of the antibiotic (Venkatesan et al., 2023). The *sul2* gene was the most prevalent gene (33.9%, 20/59) among the tested *Vibrio* isolates. This finding is in consonance with Adesiyan et al. (2022), who reported a similar *sul2* prevalence of 31.1% among the *Vibrio* species from freshwater rivers of southwest Nigeria. Despite originating from different aquatic environments, they share similar *sul2*prevalence, suggesting dissemination of *sul2* resistance genes among *Vibrio* species is not restricted to differences by habitat salinity or source of pollution. The *sul2* prevalence of this study is higher than several *Vibrio* species studies from marine aquaculture settings in China, Thailand, and Italy (Ferri et al., 2024; Jeamsripong et al., 2020; Yu et al., 2023; Zhao et al., 2018). These differences across countries and aquaculture settings suggest higher sulfonamide selection pressure in the hatchery systems, favoring proliferation of *Vibrio* isolates carrying the *sul2* gene. The *tetC* is the second highest prevalent resistant gene (28.8%, 17/59) in this study. This is lower than the findings of Sohidullah et al. (2025), who reported 60% of the *V. parahaemolyticus* carrying *tetC* gene from the shrimp ghers of Bangladesh. Shrimp is cultured in ghers for a long cycle and tetracycline is one of the most used antibiotics in ghers (Bashar et al., 2026; Kamal et al., 2015). This prolonged repeated use of tetracycline may have led to the appearance of high prevalence of *tetC* gene in shrimp ghers; however different species composition may also contribute to this discrepancy. The prevalence of the *ermB* (11.9%, 7/59) and the *strA*-*strB* (15.3%, 9/59) were comparatively lower in this study. No isolates positive to *bla*_TEM_ gene was found in this study, which is not an uncommon finding; other authors have also reported absent or very low abundance (0.8-5%) of this gene (Jeamsripong et al., 2020; Letchumanan et al., 2015; Sohidullah et al., 2025; Sony et al., 2021). The beta-lactamase-mediated resistance is common among *Vibrio* species; lower prevalence or complete absence of this gene may be mediated by intrinsic or alternative beta-lactamase genes (e.g., *bla*_CARB_-type enzymes) rather than the plasmid-borne *bla*_TEM_ gene specifically (Chiou et al., 2015; Yang et al., 2025). Association analysis between resistance genes revealed that *sul2* was significantly associated with both *strA*-*strB* and *ermB*. The *sul2* and *strA*-*strB* share a strong genetic relationship. They are often found physically linked together in a conserved multi-resistance gene cluster (*sul2*-*strA*-*strB*) on plasmids and transposons in Gram-negative bacteria, including *Vibrio* species (Okubo et al., 2019). Although no direct genetic linkage between *sul2* and *ermB* has been demonstrated, they are frequently reported by other authors to coexist within the same samples (Li et al., 2025). These co-occurrences of antibiotic resistance genes are another reason behind the appearance of MDR isolates (Algammal et al., 2025). The genetic linkage between resistance genes and mobile genetic elements raises concern about the dissemination of multi-drug resistance within the hatchery system (Okubo et al., 2019).

Aquaculture relies heavily on antibiotics to treat emerging bacterial diseases. Co-occurrence of phenotypic resistance is a major challenge in treating diseases with antibiotics, as it allows the bacteria to resist multiple drugs, leading to limited treatment options, treatment failure and emergence of super bugs (Albuquerque Costa et al., 2015; Jiang et al., 2020). In the present study, significant intra-class association was observed among the fluoroquinolones (CIP, LEV, OFX). Fluoroquinolones were also significantly associated with the quinolone nalidixic acid. This cross-resistance pattern is consistent with reports that a single quinolone resistance-determining region (QRDR) mutation in *GyrA* and *ParC* confers reduced susceptibility to quinolones and fluoroquinolones (Kherroubi et al., 2024). Significant intra-class association was also observed among the unprotected penicillins (AMP and PRL), but the penicillins with beta-lactamase inhibitor combinations (AMC and TZP) showed high susceptibility against the *Vibrio* isolates. The penicillin AMP and cephem CXM were also significantly associated. As pencillins and cehphems are beta-lactam antibiotics, they can be hydrolyzed by a shared broad-spectrum beta-lactamase (Castanheira et al., 2021). As *bla*_TEM_ was absent in the present study, this resistance may be mediated by other beta-lactamase resistance genes that were not screened in this study. Similar intra-class association was also observed among the macrolides (AZM and E), consistent with reports that mobile genetic elements carried by *Vibrio* spp. such as *mph* and *mel*/*mrx* gene clusters, confer resistance to structurally similar macrolides through efflux and drug inactivation mechanisms (Wang et al., 2018). Significant cross-class associations were observed between different antibiotics that are not structurally related, including between fluoroquinolones and tetracycline, fluoroquinolones and the macrolide azithromycin, trimethoprim-sulfamethoxazole and the macrolides/nalidixic acid, and tetracycline and nalidixic acid/ofloxacin. These antibiotics have unrelated targets and resistance mechanisms; such associations might be explained by the presence of multiple unrelated resistance genes on the same mobile genetic elements, such as plasmids and integrons. This may allow co-selection of multiple resistance genes under selection pressure from a single antibiotic (Wang et al., 2018). Both intra-class and cross-class resistance were observed in this study at comparable strength, suggesting resistance of *Vibrio* species to antibiotics is not restricted to shared mechanisms. This might add an extra layer of problem in treating diseases with antibiotics in the hatchery systems.

Association analysis between the presence of resistance genes and phenotypic resistance revealed that *sul2* was positively and significantly associated with SXT resistance. This association is consistent with established reports that *sul* genes encode an altered dihydropteroate synthase (DHPS) bearing a structural insertion that prevents sulfonamide binding, preserving normal folate metabolism (Ahmad, 2026). Significant association was also observed between the presence of *strA*-*strB* and streptomycin resistance. This association is consistent with the findings of Sunde & Norström (2005), who reported that *strA*-*strB* genes are probably involved in conferring high-level resistance to streptomycin. There were no significant associations found between the presence of the *tetC* gene and tetracycline, or between *ermB* and macrolide (AZM and E). This might be due to genes not phenotypically expressed, or resistance being mediated by other unscreened mechanisms (Kherroubi et al., 2024).

This study provides valuable information regarding *Vibrio* composition and their antibiotic resistance burden, along with associated resistance genes, across the water flow system and postlarvae of Bangladeshi shrimp hatcheries in Cox’s Bazar. Species identification was based on 16S rRNA gene sequencing of culture-dependent isolates. Future studies using next-generation sequencing-based amplicon sequencing might unravel the bacterial composition of both cultured and unculturable bacteria. Species identification based on 16S rRNA gene sequencing is not always reliable, particularly for closely related *Vibrio* species such as members of the Harveyi clade. Consequently, MLSA or WGS based studies are preferred for accurate identification and phylogenetic resolution. *Vibrio* diversity and antibiotic resistance patterns are known to vary seasonally, therefore future studies including temporal sampling are recommended. The resistance pattern across the water flow system suggested an increasing trend of antibiotic selection pressure, more samples per source would have strengthened this finding. The presence of antibiotic resistance genes pose a threat of further escalating through horizontal gene transfer (HGT). Further studies screening a broader panel of resistance genes, alongside virulence genes, would provide a comprehensive overview of the transmission potential of these traits. Frequent use of antibiotics is giving rise to antimicrobial-resistant strains, resulting in treatment failure. Alternative approaches such as probiotics, bacteriophages, and antimicrobial peptides should be prioritized for disease prevention and control.

## 5. Conclusion

This is the first study on *Vibrio* composition and antibiotic resistance patterns systematically across the hatchery water flow system and postlarvae of shrimp hatcheries in Cox’s Bazar, Bangladesh. This study also contributes to the limited body of literature on shrimp hatcheries worldwide. The findings revealed that *Vibrio* composition in the hatchery system is diverse, comprising 19 species. The Harveyi clade dominated the *Vibrio* composition and *V. alginolyticus* showed persistence across the water flow system. The *Vibrio* isolates showed high resistance to ampicillin, streptomycin, and erythromycin, and high susceptibility to chloramphenicol, nitrofurantoin, and gentamycin. The resistance pattern revealed high variability, suggesting difficulty in choosing the right antimicrobial agents for treating vibriosis outbreaks. Both the proportion of isolates with MAR index > 0.2 and MDR isolates were greater than 50%, suggesting high antibiotic selection pressure in the hatchery systems and a risk of disseminating these isolates to the ocean, farms, and the human food chain. The application of multivariable logistic regression model in predicting MDR status based on bacterial loads and resistance index in this study represents an underused approach and may offer a useful framework for future studies in aquaculture settings. Analysis revealed that the presence of *sul2* and *strA*-*strB* genes was significantly associated with their corresponding phenotypic resistance to SXT and S, respectively. Association analysis also revealed the presence of cross-class resistance among the antibiotics, suggesting that selection pressure from certain antibiotics may lead to co-selection of multiple resistance traits. Overall, this study provides crucial data on *Vibrio* composition and their resistance burden, highlighting a possible threat to human, animal, and environmental health under a One Health framework. The study can serve as a baseline for future *Vibrio* studies from marine environments and hatchery systems amid the emerging era of disease outbreaks and antibiotic treatment failure.

## Acknowledgement

We are thankful to the members of Aquatic Animal Health Group (Tahara Rohomania, Sabbir Ahmed, Shamim Ara Ripa, and Md. Ashikur Rahman Tawhid) for their generous support in laboratory work. We are also thankful to the hatchery authorities for granting permission and facilitating sample collection during this study.

## Data availability

All the 16S rRNA gene sequencing data (n = 59) generated in this study were submitted to the GenBank sequence database of the National Center for Biotechnology Information (NCBI). The accession numbers of the submitted nucleotide sequences are <u>PZ840667 to PZ840725</u>. All other data supporting the findings of this study have been included within the article or its supplemental material.

